# Unique Gluing Effect of ASXL1 K351 Monoubiquitination Stimulates PR-DUB-Mediated Nucleosomal H2AK119Ub Deubiquitination

**DOI:** 10.1101/2025.04.25.650627

**Authors:** Tianyi Zhang, Jiqing Zheng, Zebing Tong, Zhiheng Deng, Zaozhen He, Xiangwei Wu, Miao Wang, Yunxiang Du, Ziyu Xu, Shixian Tao, Xiaolin Tian, Haiteng Deng, Man Pan, Huasong Ai, Lei Liu

## Abstract

Protein ubiquitination plays a critical role in regulating fundamental biological processes through both proteolytic and nonproteolytic mechanisms. While classically known for its role in targeting proteins for degradation, ubiquitination can also modulate enzymatic activity. However, all of the known mechanisms of such modulation involve spatially constrained mechanisms that take place near the enzyme-substrate interfaces. Here, we report an unprecedented regulatory paradigm mediated by ubiquitination that is located *distal* to the enzyme-substrate interface but can still activate the polycomb repressive deubiquitinase (PR-DUB) complex. Using cryo-electron microscopy, molecular dynamics (MD) simulations, hydrogen-deuterium exchange mass spectrometry (HDX-MS), and kinetics assays, we determined that the ASXL1 K351 monoubiquitination promotes PR-DUB-mediated nucleosomal H2AK119Ub deubiquitination by stabilizing the catalytic pocket of PR-DUB, causing significant increase in maximal catalytic velocity (V_max_) but almost no change in the substrate binding affinity (K_m_). Structural analysis revealed that the ubiquitin moiety at ASXL1 K351 bridges the BAP1 and ASXL1 subunits, acting as an intramolecular “glue” that constrains the conformational dynamics of these subunits without altering the nucleosome binding interface. Further MD and HDX-MS results demonstrated the monoubiquitination of ASXL1 K351 enables PR-DUB to effectively maintain a spatial arrangement of the catalytic conformation that closely aligns with substrate cleavage. This study expands our understanding of the mechanisms of ubiquitin function to include intramolecular fastening with a molecular gluing effect and provides a deeper mechanistic understanding of PR-DUB activation in nucleosomal H2AK119Ub removal.

## Introduction

Protein ubiquitination, a versatile and multifunctional posttranslational modification, is a critical regulator that orchestrates fundamental biological processes including signal transduction, genome maintenance, and chromatin organization^1–3^. This regulatory versatility stems from its ability to dynamically modulate protein stability, protein interaction networks, and enzymatic functionality^4^. While K48-linked polyubiquitination is synonymous with proteasomal targeting^5,6^, several emerging paradigms highlight the complex role of ubiquitin in enzymatic regulation through multiple mechanistic frameworks^7^. First, as a principal mechanism, ubiquitination drives intramolecular recruitment via ubiquitin recognition domains (e.g., UIM, UBA, CUE, and BUZ)^8,9^, where the ubiquitin modification serves as a molecular scaffold to increase substrate engagement and catalytic efficiency. As a typical example, the PRC2 accessory subunits Jarid2 and Aebp2 synergistically recognize H2AK119Ub-modified nucleosomes through their UIM and C2H2 zinc finger domains, respectively, to potentiate the H3K27 methyltransferase activity of the PRC2.2 complex^10,11^. Second, as a complementary mechanism, substrate ubiquitination-induced conformational gating can occur, in which substrate ubiquitination restricts enzyme conformation dynamics through steric constraints to favor catalytically active states, as exemplified in the activation process of the H3K79 methyltransferase Dot1L by H2BK120Ub^12–14^. Furthermore, our recent work revealed a third regulatory mode involving ubiquitin-dependent structural dynamics, in the case of Dot1L-mediated H3K79 methylation, which is activated allosterically by H2BK34Ub-induced nucleosome reshaping^15^. Notably, the above three models share a common spatial feature, where ubiquitination event occurs *near* the enzyme-substrate interface to modulate binding affinity or catalytic competency.

Here we report an unprecedented *distal* ubiquitin regulatory paradigm distinct from the three aforementioned mechanisms, which was serendipitously discovered in our investigation of the PR-DUB complex. This complex, which is composed of a catalytic subunit BAP1 and one of the ASXL family auxiliary subunits (ASXL1/2/3), selectively catalyzes the removal of nucleosomal monoubiquitination at histone H2A lysine 119 (H2AK119Ub)^16^. This deubiquitination activity establishes a dynamic equilibrium with PRC1-mediated ubiquitination, maintaining the H2AK119Ub levels required for polycomb-dependent transcriptional repression^17–20^. Oncogenic mutations in BAP1/ASXL, particularly at residues surrounding the catalytic pocket or substrate-binding interfaces, implicate PR-DUB dysregulation as a potential oncogenic driver^21–27^. BAP1 displays low intrinsic catalytic activity, but its interaction with the DEUBAD domain of ASXL proteins stimulates deubiquitination activity by improved substrate ubiquitin recognition and BAP1/ASXL complex stabilization^28–30^. Intriguingly, conserved lysine residues within DEUBAD domains of ASXL proteins (ASXL1 K351, ASXL2 K370, and ASXL3 K350) have been identified as ubiquitination sites through proteomics profiling reported in the PhosphoSitePlus database^31–33^, with cellular studies consistently demonstrating their predominant monoubiquitination status^29^. Recent biochemical and functional studies have revealed that ASXL monoubiquitination stabilizes ASXL proteins, augments PR-DUB catalytic activity, and regulates cell proliferation and development processes^29^. Nevertheless, the structural mechanisms by which ASXL monoubiquitination potentiates PR-DUB-mediated H2AK119Ub removal remain poorly understood.

In this study, we applied an integrated multidisciplinary approach combining cryo-electron microscopy (cryo-EM), molecular dynamics (MD) simulations, hydrogen-deuterium exchange mass spectrometry (HDX-MS), and mutational analyses with functional deubiquitination assays, delineating a noncanonical mechanism of allosteric PR-DUB activation mediated by ASXL1 K351 monoubiquitination. Our findings reveal that this ubiquitin modification that locates *distal* to the enzyme-substrate interface serves as a glue between the BAP1 and ASXL1 subunits and thereby constrains the conformational dynamics of the PR-DUB complex and stabilizes the deubiquitinating catalytic pocket, while not alters either the intensity or spatial configuration of enzyme-substrate interactions. This mechanistic paradigm deviates fundamentally from the canonical view of ubiquitination as a canonical signalling or degradation-promoting moiety that typically functions at substrate-proximal enzyme-substrate interfaces. Thus, this work expands our understanding of the versatile regulatory repertoire of ubiquitin as an intramolecular fastener with a molecular gluing effect and provides a deeper mechanistic understanding of the PR-DUB in nucleosomal H2AK119Ub removal.

## Results

### ASXL1 K351 monoubiquitinated PR-DUB stimulates the nucleosomal H2AK119 deubiquitination activity

Our study was initiated with the preparation of ASXL1 K351 ubiquitinated PR-DUB. The UBE2E family of E2 ubiquitin-conjugating enzymes has been demonstrated to mediate the E3 ligase-independent site-specific monoubiquitination of ASXL family proteins^29^. Our previously engineered UBE2E1* variant demonstrated significantly enhanced substrate ubiquitination activity^34^. Thus, we applied the UBE2E1*-based chemoenzymatic approach to generate monoubiquitinated BAP1/ASXL1 complex. Full-length BAP1 (residues 1–729) was coexpressed with the ASXL1 DEUBAD domain (residues 238–390) in *Escherichia coli*, followed by purification of the reconstituted complex (**Extended Data Fig. 1a, b**). A subsequent *in vitro* ubiquitination experiment was conducted using a system containing Uba1 (E1), our engineered UBE2E1* (E2), ubiquitin, the BAP1/ASXL1 complex and ATP. Size-exclusion chromatography and SDS–PAGE analysis confirmed the successful generation of the monoubiquitinated BAP1/ASXL1 complex (**Fig. 1a** and **Extended Data Fig. 1c-e**). Mass photometric analysis revealed a molecular weight increase of ∼7 kDa (102 kDa *vs.* 109 kDa) for monoubiquitinated BAP1/ASXL1 complex relative to its unmodified counterpart, consistent with the covalent attachment of a single ubiquitin moiety (**Extended Data Fig. 1f-g**). The specificity of ubiquitination at K351 was rigorously validated through multiple experiments: (1) abrogation of ubiquitination in ASXL1 K351R mutants (**Fig. 1b**) and (2) tandem mass spectrometry (MS/MS) identification of ubiquitinated peptides spanning K351 (**Fig. 1c**).

**Figure 1.**
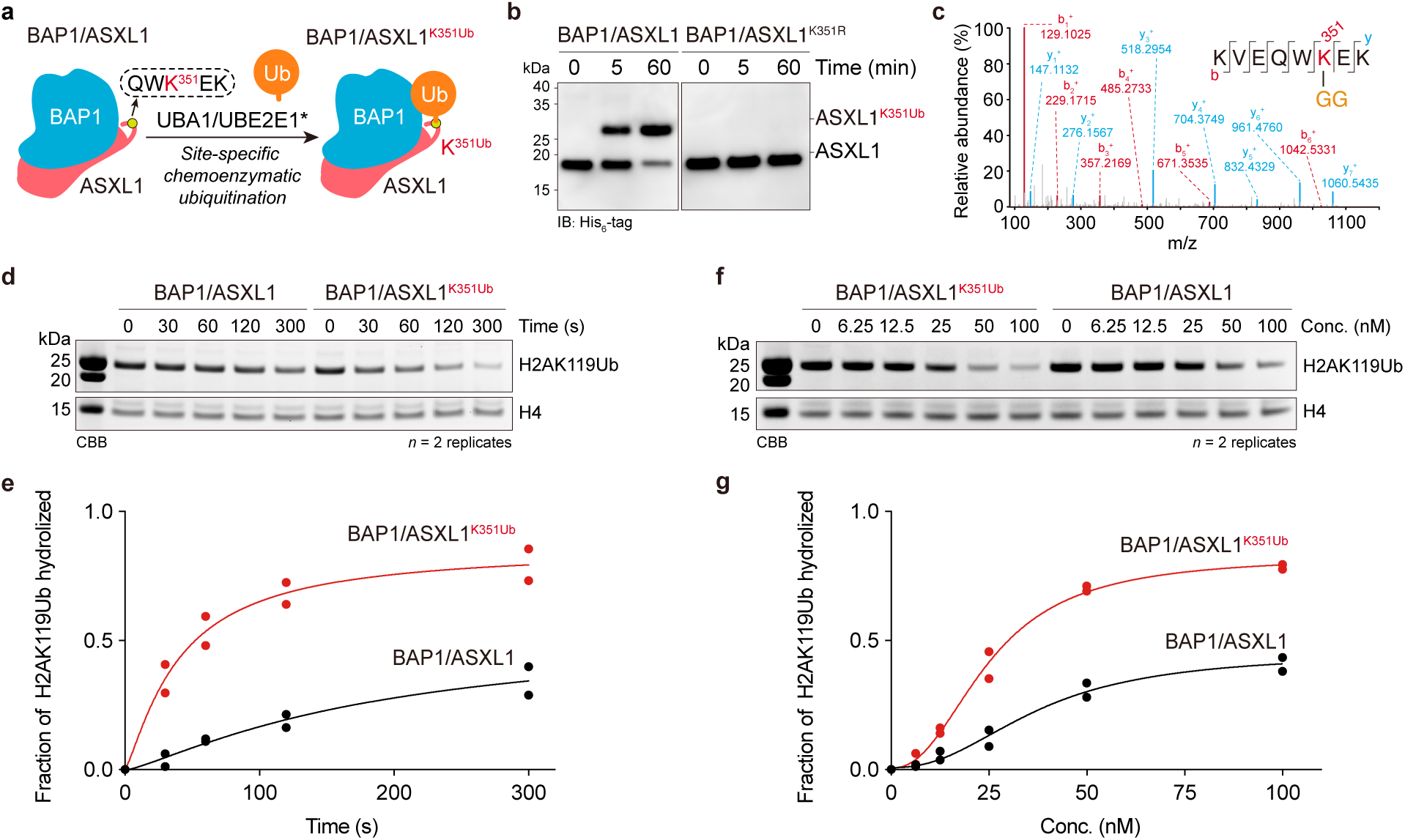
Monoubiquitination of ASXL1 K351 enhances PR-DUB activity on H2AK119Ub nucleosomes. **a**, Schematic representation of UBE2E1-mediated site-specific monoubiquitination at ASXL1 lysine 351 (K351). **b, c,** UBE2E1 catalyzes site-specific monoubiquitination of ASXL1. **b,** Western-blot analysis of UBE2E1-mediated ubiquitination of BAP1/ASXL1 complexes. BAP1/ASXL1 (left) or BAP1/ASXL1-K351R (right) was incubated for 0, 5 or 60 min. Labeled proteins were analyzed by immunoblotting using an anti-His_6_ antibody. **c,** Tandem mass spectrometry identifies ASXL1-Ub as the ubiquitination product and confirms K351 as the modification site. **d, e,** Time-course analysis reveals enhanced catalytic activity of BAP1/ASXL1^K351Ub^ compared to BAP1/ASXL1 in deubiquitinating H2AK119Ub nucleosomes. **f, g,** Concentration-dependent analysis demonstrates a higher activity on H2AK119Ub nucleosome of BAP1/ASXL1^K351Ub^ than BAP1/ASXL1. Gel images shown in **d** and **f** are representative of independent biological replicates (n=2).

Next, we conducted comparative nucleosomal deubiquitination experiments to assess the effect of ASXL1 K351 monoubiquitination on PR-DUB activity. Using chemically synthesized H2AK119Ub nucleosomes containing native isopeptide bonds as substrates (800 nM), we carried out nucleosomal deubiquitination assays with enzyme concentrations of 50 nM in reaction buffer (50 mM HEPES, 150 mM NaCl, and 1 mM DTT). Time-course analysis revealed progressive attenuation of H2Aub band intensity, with notably accelerated decay kinetics observed when using ASXL1 K351 monoubiquitinated PR-DUB (hereafter BAP1/ASXL1^K351Ub^) compared with the non-ubiquitinated PR-DUB complex (**Fig. 1d-e**). Subsequent titration of deubiquitinase concentrations (6.25–100 nM) under fixed substrate conditions (800 nM) consistently increased the catalytic efficiency of Ub-PR-DUB by approximately 2.0∼3.0-fold increase, as evidenced by more rapid elimination of the H2Aub signal at equivalent enzyme concentrations (**Fig. 1f-g**). This finding is consistent with previous qualitative analysis using Flag-H2A purified native nucleosomes^29^. Collectively, these biochemical results establish that ASXL1 K351 monoubiquitination stimulates PR-DUB-mediated deubiquitination of H2AK119Ub nucleosomes.

### Capturing the structure of the Ub-PR-DUB-mediated nucleosomal H2AK119Ub deubiquitination using activity-based probes

To investigate the mechanism by which ASXL1 K351 monoubiquitination enhances PR-DUB activity, we performed the cryo-EM to resolve the structure of BAP1/ASXL1^K351Ub^ bound to H2AK119Ub nucleosomes. We developed an activity-based deubiquitinase trapping strategy that enables covalent capture of the transient catalytic intermediate during BAP1-mediated nucleosome deubiquitination (**Fig. 2a**). The H2AK119Ub^AT^ nucleosome features a strategically positioned disulfide-activated thiol moiety adjacent to the isopeptide bond, enabling proximity-driven conjugation with the catalytic cysteine in the PR-DUB active site. We first synthesized the histone H2AK119Ub^AT^ (the synthetic routine and characterization were provided in **Extended Data Fig. 2**), and were then assembled into octamers with H2B, H3 C96S/C110S (mutations introduced to prevent intra-histone disulfide exchange) and H4, followed by reconstitution with the 147-bp 601 positioning DNA to assemble nucleosome core particles (NCPs). The incubation of this activity-based nucleosomal deubiquitinase probe with BAP1/ASXL1^K351Ub^ resulted in the covalent crosslinking of BAP1 and H2Aub, generating a stable structural mimic of the tetrahedral hydrolytic transition state (**Fig. 2d-e**). ASXL1 K351 monoubiquitination enhances the crosslinking efficiency of PR-DUB for H2AK119Ub^AT^ nucleosomes, as evidenced by accelerated depletion of both the H2Aub and BAP1 bands (**Fig. 2b-c**).

**Figure 2.**
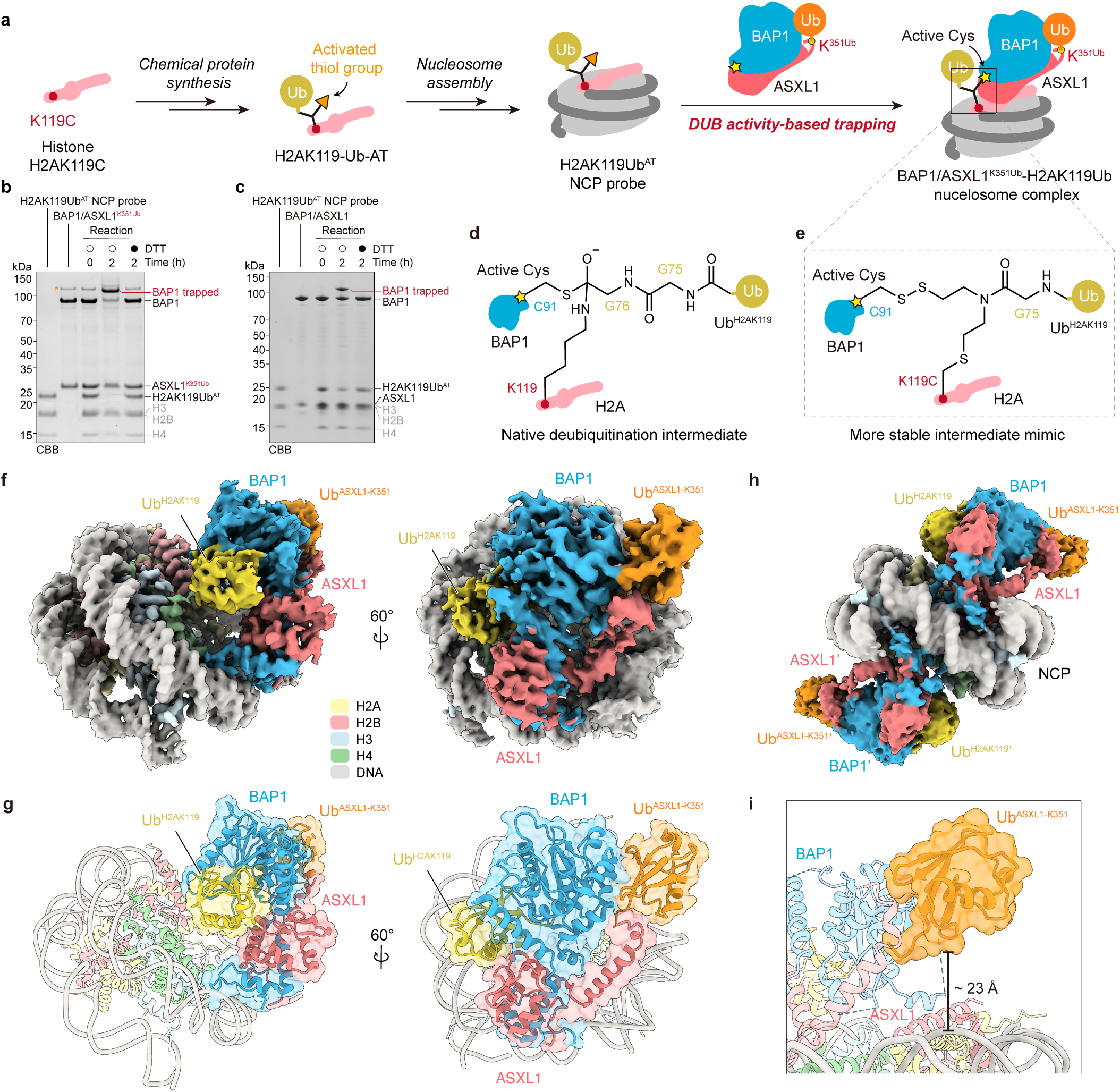
Structural insights into BAP1/ASXL1^K351Ub^-mediated H2AK119Ub nucleosome deubiquitination captured using activity-based probe. **a**, Schematic representation of activity-based probe trapping strategy. **b, c,** Coomassie-stained SDS-PAGE analysis of H2AK119Ub^AT^ nucleosome probe crosslinked with BAP1/ASXL1 or BAP1/ASXL1^K351Ub^. Time-dependent crosslinking (0- and 2-hour time points) of the BAP1/ASXL1^K351Ub^ or BAP1/ASXL1 complex to H2AK119Ub nucleosomes under reducing (+DTT) and non-reducing (–DTT) conditions, resolved by SDS-PAGE. The asterisk-denoted band corresponds to the intact His-SUMO-BAP1, with molecular weight markers indicated on the left (kDa). **d,** Schematic of the native transition state of H2AK119Ub nucleosome deubiquitination by BAP1/ASXL1. BAP1 acts as the catalytic subunit in the BAP1/ASXL1 complex. **e,** Schematic representation of designed intermediate structure using the activity-based probe, which highly mimics the native transition state shown in **d**. **f,** Cryo-EM density of the 1:1 BAP1/ASXL1^K351Ub^/H2AK119Ub nucleosome complex. **g,** Atomic model of the 1:1 complex. **h,** Cryo-EM density of the 2:1 BAP1/ASXL1^K351Ub^/H2AK119Ub nucleosome complex. **i,** Close-up view of the spatial relationship between the Ub^ASXL1-K351^ and the nucleosome, the Ub^ASXL1-K351^ is positioned about 23 Å above the nucleosomal surface (distance measured between the α-carbon atom of Ub^ASXL1-K351^ D39 and C4’ atom of DNA DA-62).

We subjected the covalently stabilized complex of BAP1/ASXL1^K351Ub^ and H2AK119Ub nucleosome to cryo-EM structural analysis. A total of 7994 micrographs were collected using a 300 kV Titan Krios microscope and processed through RELION 3.1, yielding two reconstructions: a 3.12 Å resolution structure of the PR-DUB complex bound to the H2AK119Ub nucleosome in 1:1 stoichiometry and a 3.27 Å resolution structure with binding in 2:1 stoichiometry (**Fig. 2f-h, Extended Data Fig. 3-4** and **Extended Data Table 1**). As both cryo-EM reconstructions exhibited a similar organization, we focused our subsequent analysis on the higher-resolution 1:1 complex. In this structure, the ubiquitin modification on the substrate H2AK119 (Ub^H2AK1^^19^) was encapsulated by a bipartite interface formed by BAP1 and ASXL1 (**Fig. 2g**). The ASXL1 K351-modified ubiquitin moiety (Ub^ASXL1-K3^^51^) resides at the C-terminal periphery of ASXL1 (residues 247–357) and interacts with residues 110–133 of the BAP1 UCH domain. This Ub^ASXL1-K3^^51^ adopts a unique spatial configuration, positioned approximately 23 Å above the nucleosomal surface on the dorsal aspect of the PR-DUB substrate-binding interface, with no direct contact observed for Ub^H2AK1^^19^ and the nucleosome substrate (**Fig. 2i**).

### ASXL1 K351 monoubiquitination does not alter the binding mode between the PR-DUB and nucleosomes

We performed a comparative structural analysis between our resolved structure of BAP1/ASXL^K351Ub^/H2AK119Ub nucleosome and the previously reported complex structure of BAP1/ASXL1 bound to the H2AK119Ub chromatosome (PDB: 8H1T)^35^. The overall architectures demonstrated high similarity with a root mean square deviation (RMSD) of 0.917 Å for all aligned atoms, except that ASXL1 K351Ub was present in one structure and absent in the other (**Fig. 3a-b**). We also extended this comparative analysis to include another published BAP1/ASXL1/H2AK119Ub nucleosome complex (PDB: 8SVF)^36^, revealing comparable structural features (**Extended Data Fig. 5a-b**). Detailed analysis of the PR-DUB/nucleosome interface revealed conserved interaction networks across four critical regions: (1) the C-terminal extension (CET) domain of BAP1 (residues 699–715) engaging nucleosomal DNA SHL0, (2) the α-helical segment (residues 56–60) of BAP1 anchoring into the H2A–H2B acidic patch, (3) the catalytic triad (C91/H169/D184) within the BAP1 deubiquitinase active site, and (4) the BAP1/ASXL1 heterodimerization interface (**Fig. 3c-f**). Notably, both the residue types and interaction geometries at these interfaces remained unchanged, with conservation of the polar contacts and hydrophobic packing interactions.

**Figure 3.**
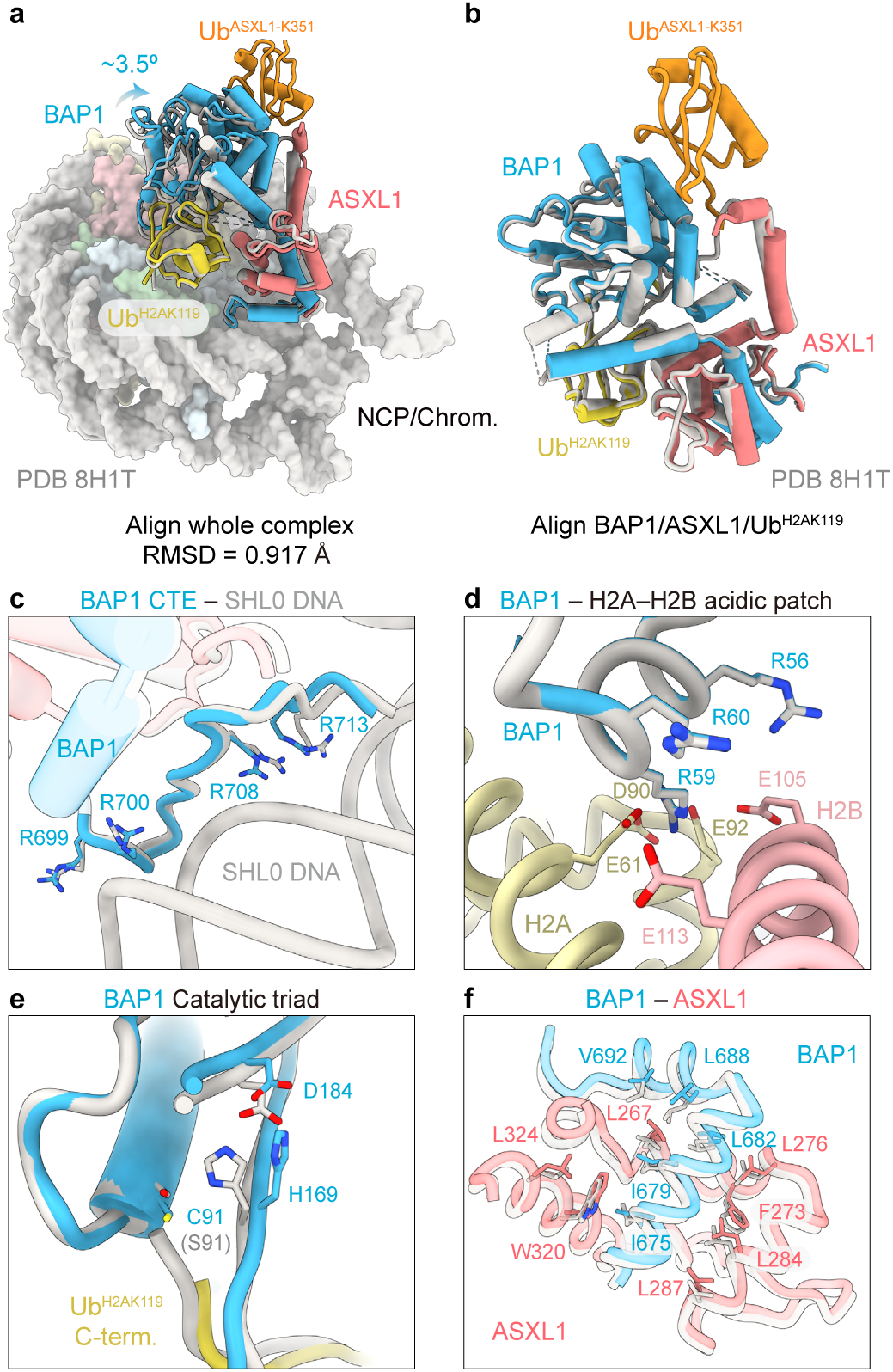
ASXL1 monoubiquitination does not alter the binding mode between the PR-DUB and the nucleosome. **a, b**, Structural superposition of the BAP1/ASXL1^K351Ub^/H2AK119Ub nucleosome complex with the BAP1/ASXL1/H2AK119Ub chromatosome structure (PDB: 8H1T), revealing no significant conformational difference. **a,** Global alignment of the two complexes (RMSD = 0.917 Å). **b,** Alignment of the BAP1/ASXL1/Ub^H2AK119^ (RMSD = 1.029 Å). **c-f,** Close-up view of key interfaces of PR-DUB and nucleosomes. **c,** The BAP1 C-terminal extension (CTE) domain interacts with SHL0 DNA interface. **d,** BAP1 residues 56-60 form an α-helix binding to H2A-H2B acidic patch. **e,** Spatial arrangement of BAP1 catalytic triad (C91, H169, D184). **f,** BAP1-ASXL1 heterodimerization interface.

Furthermore, we performed amplified luminescent proximity homogeneous assays (AlphaLISA) and electrophoretic mobility shift assays (EMSAs) to investigate the impact of ASXL1 K351 monoubiquitination on the binding affinity of PR-DUB for H2AK119Ub nucleosomes. A catalytically inactive BAP1 mutant (C91A) was used to prevent H2AK119 nucleosome cleavage by PR-DUB. The overlapping binding curves observed from AlphaLISA (**Extended Data Fig. 5c**) and the comparable band shift patterns in EMSA (**Extended Data Fig. 5d-e**) convincingly demonstrated that monoubiquitination at ASXL1 K351 does not alter the nucleosome-binding capacity of PR-DUB. These structural and biochemical results suggest that ASXL1 K351 monoubiquitination neither alters the intrinsic nucleosome-binding mode or strength of PR-DUB, nor creates *de novo* interfacial contacts between PR-DUB and nucleosomal substrates.

### Ub^ASXL1-K351^ is sandwiched between BAP1 and ASXL1

We next performed detailed structural analysis focusing on the ubiquitin moiety conjugated at the ASXL1 K351 site. Ub^ASXL1-K351^ was intermolecularly clamped between the C-terminus of the ASXL1 DEUABD, making contact with the helix (residues 110–133) of the BAP1 UCH domain (**Fig. 4a**). Ub^ASXL1-K351^ was structurally positioned at the interface between the BAP1 and ASXL1 subunits, forming a sandwiched configuration (**Fig. 4a**). Specifically, the C-terminus of ubiquitin at the L71/L73 patch and the L8/T9 loop interact with BAP1 residues N133 and R114/F118, respectively (**Fig. 4b-c**). The I36 patch of ubiquitin was positioned adjacent to ASXL1 residues 353– 354 (**Fig. 4d**). Mutations at the canonical Ub-interacting sites on BAP1 to alanine attenuated the stimulating effect of ASXL1 K351 monoubiquitination on nucleosomal H2AK119Ub deubiquitination (**Fig. 4e** and **Supplementary Fig. 1**). Moreover, comparative cryo-EM analysis revealed that the C-terminal helix (residues 340–357) of the ASXL1 DEUABD domain was considerably more stable and ordered in our structure than in previous ASXL1 K351 non-ubiquitinated reconstitutions (EMD-35180^35^ and EMD-40789^36^) (**Extended Data Fig. 5f-h**). These results indicate that Ub^ASXL1-K351^ plays a crucial role in bridging the two subunits of PR-DUB, enhancing the stability of the ASXL1 DEUBAD tail helix and increasing the deubiquitination activity of PR-DUB towards the H2AK119Ub nucleosome.

**Figure 4.**
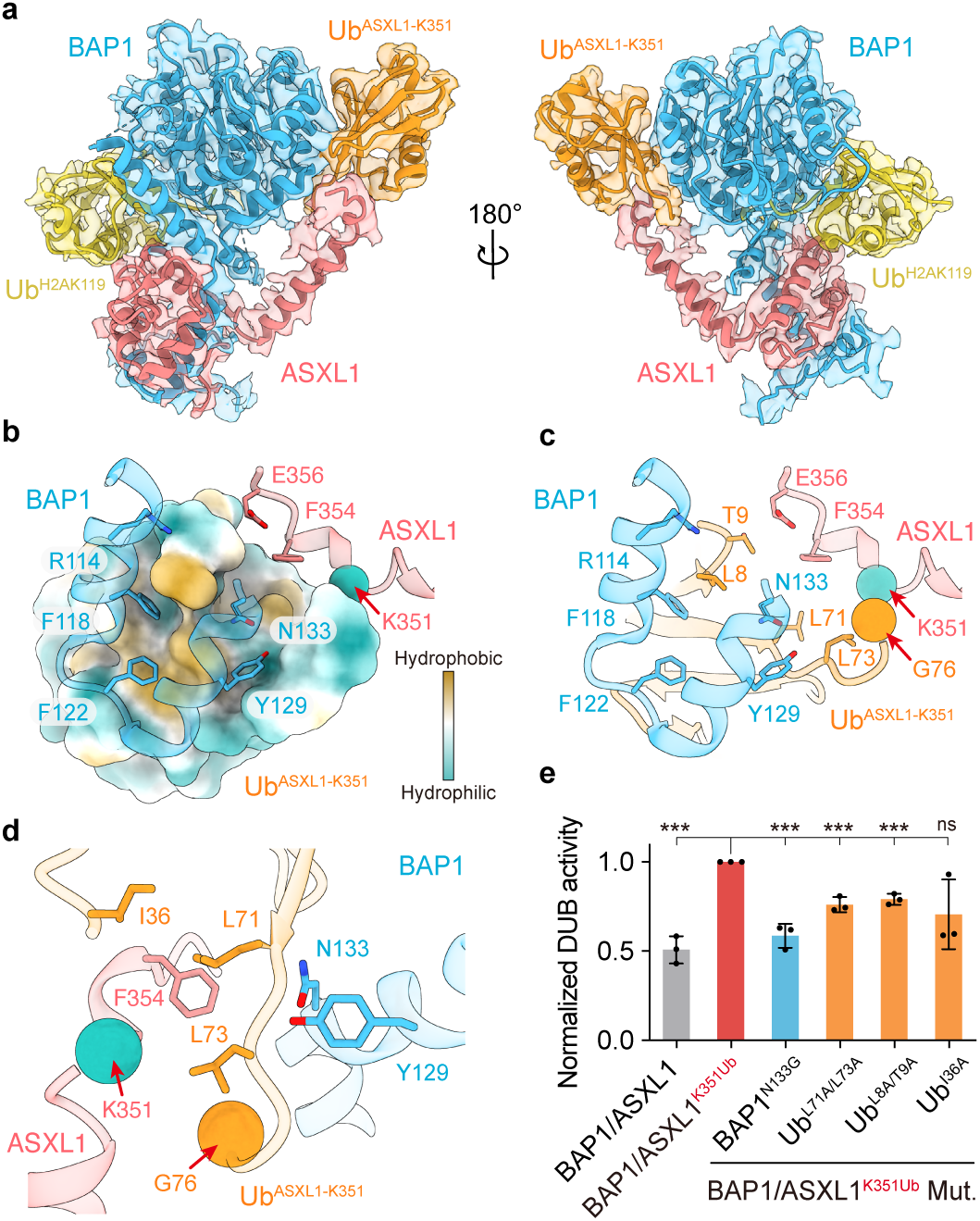
Ub^ASXL1-K351^ mediates inter-subunit interactions of BAP1 and ASXL1 to augment catalytic activity toward H2AK119Ub nucleosomes. **a**, Structural representation of BAP1/ASXL1^K351Ub^ bound to the substrate Ub^H2AK119^, with the corresponding cryo-EM density transparency embed. **b, c** Close-up view of the interaction of the BAP1 and ASXL1 segment around K351. **d,** Close-up atomic view of Ub^ASXL1-K351^ and BAP1/ASXL1 interaction. **e,** Quantified activity of BAP1/ASXL1^K351Ub^ and BAP1/ASXL1^K351Ub^ mutants (BAP1-N133G, Ub-L71A/L73A, Ub-L8A/T9A, Ub-I36A), and WT BAP1/ASXL1. Data show mean ± s.d. from three independent experiments.

### ASXL1 K351 monoubiquitination stimulates the intrinsic deubiquitination activity of PR-DUB

The above structural and biochemical analyses demonstrated that ASXL1 K351 monoubiquitination neither interacts with nucleosomal substrates nor alters the binding modality of the PR-DUB complex for nucleosomes. We subsequently compared the deubiquitinase activity of PR-DUB on two non-nucleosomal substrates: Ub-AMC (Ubiquitin-7-amido-4-methylcoumarin) and a fluorescently labelled K119-ubiquitinated H2A(86–129) peptide (chemical synthetic route and characterization details in **Extended Data Fig. 6a-e**). Remarkably, time-course deubiquitination assays revealed that, compared with non-ubiquitinated BAP1/ASXL1, BAP1/ASXL1^K351Ub^ exhibited considerable enhanced activity towards both ubiquitinated H2A peptide and Ub-AMC substrates (**Extended Data Fig. 6f-g**). To quantify this activation, we conducted Michaelis‒Menten kinetic analysis under steady-state conditions (100 nM enzyme concentration, 2.5–40 μM substrate gradient). The Michaelis constant (K_m_) values of BAP1/ASXL1 and BAP1/ASXL1^K351Ub^ were comparable for the K119-ubiquitinated H2A(86–129) peptide (9.7 ± 1.9 μM vs. 8.9 ± 1.4 μM) or Ub-AMC (18.8 ± 3.8 μM vs. 15.1± 3.6 μM) (**Fig. 5a-b**). This conservation of Kₘ values aligned with the similar substrate binding capacities for nucleosomal substrates observed in the previous AlphaLISA and EMSA experiments (**Extended Data Fig. 5c-e**). In contrast, the maximal velocity (V_max_) was markedly increased for K351 ubiquitinated ASXL1 for both the ubiquitinated H2A peptide (0.03 ± 0.002 μM/s vs. 0.09 ± 0.006 μM/s) and Ub-AMC (0.03 ± 0.003 μM/s vs. 0.08 ± 0.008 μM/s) (**Fig. 5a-b**). Collectively, these results suggest that K351 monoubiquitination of ASXL1 enhances the catalytic efficiency of the PR-DUB complex across diverse substrates including ubiquitinated nucleosomes or ubiquitinated peptides, and this increased activity stems not from increased enzyme-substrate binding affinity but rather from an increase in V_max_.

**Figure 5.**
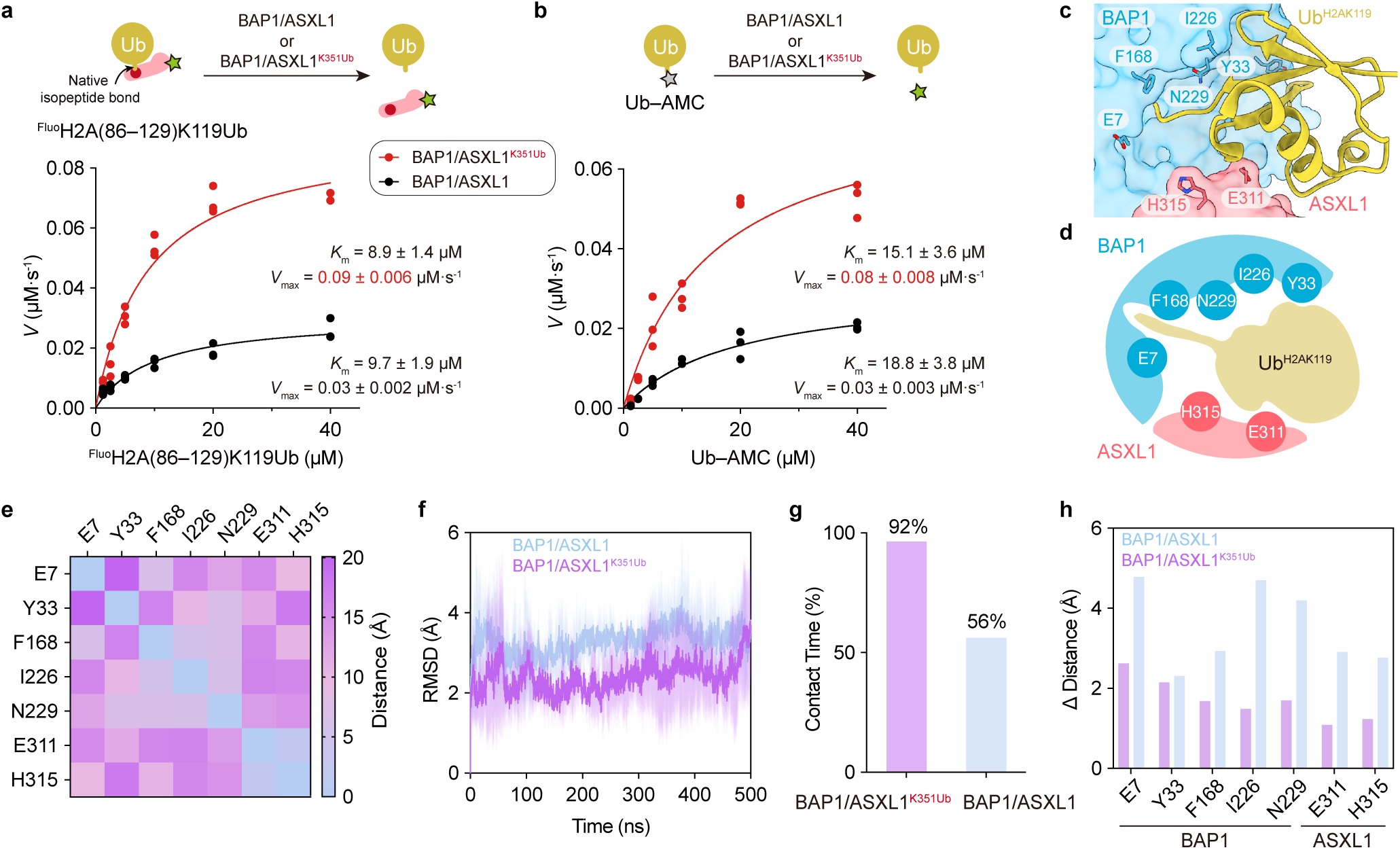
ASXL1 K351 monoubiquitination stimulates the PR-DUB intrinsic deubiquitination activity through stabilization of the catalytic pocket of PR-DUB. **a, b**, Michaelis-Menten kinetics for BAP1/ASXL1 or BAP1/ASXL1^K351Ub^ catalyzing deubiquitination of fluorescent-H2A (86-129)-K119Ub peptide **(a)** or Ub-AMC **(b)**. Data represent mean ± s.d. (n = 3). **c,** Structural representation of the BAP1/ASXL1 catalytic pocket, seven key residues (BAP1: E7, Y33, F168, I226, N229; ASXL1: E311, H315) interacting with substrate Ub (yellow) are highlighted. **d,** A schematic cross-sectional view of the catalytic pocket, with key residues (circles) involved in substrate Ub^H2AK119^ coordination. **e,** Heatmap of the minimum pairwise distances between the key residues of the initial structure used in MD analysis. **f,** RMSD of the key residues over the 500 ns MD simulation trajectory (BAP1/ASXL1^K351Ub^: pink; BAP1/ASXL1: light blue). **g,** Statistical analysis of contact time for key residues. Contact time: the percentage of total simulation time during which all pairwise distances between five critical residues on BAP1 (E7, Y33, F168, I226, N229) and two critical residues on ASXL1 (E311, H315) remain below 20 Å. **h,** α-carbon distance comparison between the initial structure and the most populated MD cluster structure for the seven key residues.

### ASXL1 K351 monoubiquitination increases the stability of the PR-DUB catalytic pocket

To understand the mechanism by which ASXL1 K351 monoubiquitination stimulates the V_max_ of PR-DUB, we conducted comparative molecular dynamics (MD) analyses of the BAP1/ASXL1^K351Ub^ and BAP1/ASXL1 complexes. All-atom MD simulations were performed under periodic boundary conditions using the OPLS4 force field and TIP3P explicit water model, with three independent 500-ns production runs for each system. Initial velocity distributions were generated using distinct random seeds to ensure comprehensive sampling of the conformational space. We focused on our analysis on the correlation of the structural rigidity and integrity of the substrate ubiquitin-binding pocket with the deubiquitination activity of PR-DUB. Specifically, we used 500-ns trajectory analysis to examine the structural deviations of seven critical residues (BAP1 residues E7, Y33, F168, I226 and N229; ASXL1 residues E311 and H315), which collectively create the recognition pocket encapsulating the substrate ubiquitin (**Fig. 5c-e**). The average RMSD of BAP1/ASXL1^K351Ub^ was 2.39 Å, which was lower than the 3.29 Å observed in BAP1/ASXL1 (**Fig. 5f**). Furthermore, BAP1/ASXL1^K351Ub^ maintained a complete pocket conformation for 92% of the simulation time, exceeding the 56% observed in non-ubiquitinated BAP1/ASXL1 (**Fig. 5g**). Clustering analysis of the simulation trajectories revealed that the RMSD of the predominant cluster structure (largest cluster) of BAP1/ASXL1^K351Ub^ complex relative to the initial conformation was 1.71 Å, which was significantly lower than the 3.52 Å of the corresponding cluster in the BAP1/ASXL1 complex (**Fig. 5h** and **Extended Data Fig. 7a-d**). Similar trends were observed in the secondary cluster structure (second largest cluster) (**Extended Data Fig. 7e-f** and **Supplementary Fig. 2**). These MD data suggest that monoubiquitination at ASXL1 K351 enables PR-DUB to effectively maintain a spatial arrangement of the catalytic conformation that closely aligns with substrate cleavage, thereby facilitating a highly efficient catalytic reaction.

To complement the MD simulations, we performed hydrogen-deuterium exchange mass spectrometry (HDX-MS) to probe the impact of ASXL1 K351 monoubiquitination on the conformational dynamics of the PR-DUB complex. Both BAP1/ASXL1 and BAP1/ASXL1^K351Ub^ complexes were deuterium labelled at four time points (30 s, 90 s, 300 s, and 1800 s), followed by tryptic digestion and high-resolution mass spectrometry (**Extended Data Fig. 8a**). Peptide-level deuterium uptake maps were systematically constructed for comparative analysis of the conformational dynamics of ubiquitinated and non-ubiquitinated PR-DUB (**Extended Data Fig. 8b-c** and **Supplementary Fig. 3**). Regions proximal to Ub^ASXL1-K351^ — specifically, BAP1 (113–128) and ASXL1 (330–354) — exhibited reduced deuterium exchange rates in the BAP1/ASXL1^K351Ub^ complex (**Extended Data** Fig. 10d–g). This decreased solvent accessibility corroborated our structural data showing that K351 monoubiquitination glues BAP1 to the ASXL1 C-terminal helix (**Fig. 4a**). Remarkably, hydrogen–deuterium exchange analysis revealed reduced deuterium uptake in the ASXL1 (313–334) segment of the BAP1/ASXL1^K351Ub^ complex compared with its non-ubiquitinated counterpart (**Extended Data Fig. 8h-i**). The ASXL1 (313–334) segment constitutes a critical helical region that encapsulates the substrate ubiquitin, suggesting that monoubiquitination at ASXL1 K351 enhances the structural stability of this functionally essential element around the catalytic pocket.

## Discussion

This study elucidates the structural and mechanistic basis through which ASXL1 K351 monoubiquitination augments PR-DUB-mediated deubiquitination of H2AK119Ub nucleosomes. Through a combination of biochemical assays, cryo-EM structural analysis, kinetic profiling, MD simulations and HDX-MS, we demonstrated that ASXL1 K351Ub allosterically promotes the PR-DUB activity reflected in enhanced V_max_ that may result from the stabilization of the deubiquitinating catalytic pocket. Unlike established ubiquitination regulatory paradigms — such as substrate-proximal recruitment (e.g., PRC2 recruitment by H2AK119Ub)^10^, conformational gating (e.g., Dot1L activation by H2BK120Ub)^12^, or substrate structure reshaping (e.g., H2BK34Ub-induced nucleosome restructuring augments Dot1L activity)^15^ — ASXL1 K351Ub operates through a unique distal allosteric axis. This ubiquitin moiety bridges the BAP1 UCH domain (residues 110–133) and the ASXL1 DEUBAD helix (residues 313–334), stabilizing a conformationally dynamic catalytic pocket while preserving native substrate-binding interfaces. Quantitative kinetic analyses revealed that ASXL1 K351 monoubiquitination increases the V_max_ by 2.0- to 3.0-fold across both nucleosomal and non-nucleosomal substrates without altering its K_m_ value, indicating enhanced catalytic turnover rather than improved substrate affinity. The MD trajectories and HDX-MS deuterium uptake maps converge to demonstrate that this kinetic acceleration arises from the reduced conformational dynamics in the substrate ubiquitin-binding pocket. Ub^ASXL1-K351^-mediated inter-subunit clamping restrains fluctuations effectively, facilitating the pre-organization of the active site for efficient hydrolysis of the isopeptide bond (**Fig. 6**). The identification of this unique ubiquitin-mediated gluing effect expands the functional repertoire of ubiquitination beyond classical signalling or degrading-related roles, establishing it as a structural adhesive mechanism capable of allosterically reprogramming enzyme complex architecture.

**Figure 6.**
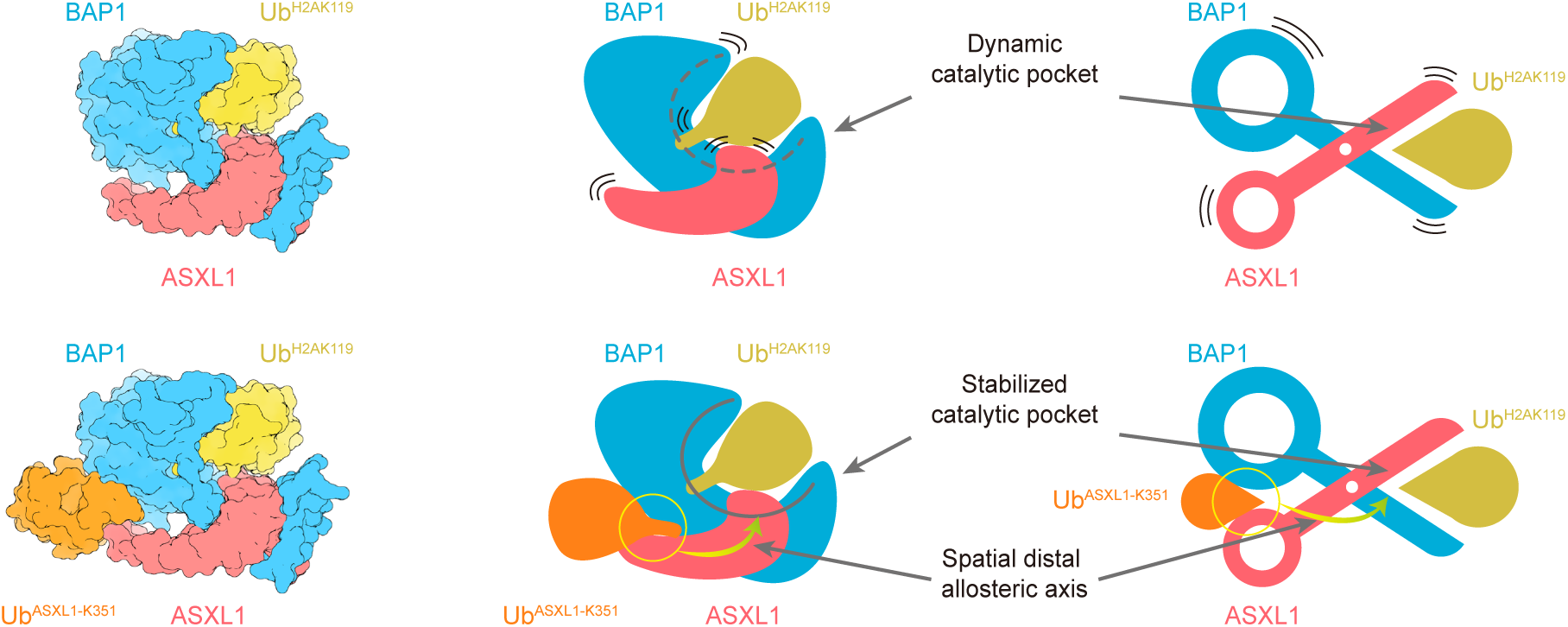
Mechanistic model of ASXL1 K351 monoubiquitination mediated PR-DUB activity activation. The PR-DUB complex exhibits a dynamic scissor-like architecture with two functional states: (1) In the unmodified state (absence of ASXL1-K351 monoubiquitination), the BAP1/ASXL1 heterodimer displays inherent conformational flexibility that permits scissor-like motions, resulting in suboptimal alignment of catalytic residues. (2) Conversely, ASXL1-K351 monoubiquitination triggers ubiquitin-induced long-range allosteric stabilization of the scissor, positioning the PR-DUB catalytic pocket ready for the isopeptide bond cleavage.

Our structural and biochemical analyses demonstrated that, in contrast to synthetic small molecular glues such as cereblon E3 ligase modulatory drugs (CELMoDs), which create neo-interfaces for targeted degradation, Ub^ASXL1-K351^ reinforces preexisting intersubunit contacts in the PR-DUB complex, representing an evolutionarily optimized proteinaceous glue. This unveils a nonclassical regulatory paradigm in which ubiquitin functions as an intramolecular fastener that stabilizes a specific conformation, extending its functional spectrum beyond conventional signalling to orchestrate macromolecular assembly by modulating intramolecular mechanics. Our observations align with other emerging evidence of ubiquitin’s nonproteolytic structural roles in stabilizing complexes^7,37^, for example in the Fanconi anemia (FA) pathway, where monoubiquitination of Fanconi anemia D2 (FANCD2) at K561 enhances the binding affinity of the ID2 complex to dsDNA at stalled replication forks or DNA crosslink lesions^38,39^. Strikingly, the BAP1-Ub^ASXL1-K3^^51^ interface achieves stabilization through an unconventional bipartite interaction with the residues R114 and N113 in BAP1 and the residues 353–354 in ASXL1. This dual anchoring strategy bypasses classical ubiquitin-binding domains such as UBA or UIM. This finding suggests that the intrinsic plasticity of ubiquitin could enable its repurposing as a versatile glue across diverse biological contexts, independent of canonical recognition motifs.

The BAP1-Ub^ASXL1-K351^ interface has emerged as a mechanistically distinct vulnerability in malignancies driven by PR-DUB dysregulation, particularly BAP1-deficient mesotheliomas and ASXL1-mutant myeloid neoplasms^23,27^. Our structural delineation of this ubiquitin-mediated gluing interface provides the structural basis for developing therapeutic modalities that target PR-DUB hyperactivity. Structure-guided design of small-molecule interfacial inhibitors or constrained peptidomimetics mimicking the BAP1-Ub^ASXL1-K351^ interaction surface might disrupt the allosteric activation mechanism. Alternatively, pharmacological inhibition of the UBE2E1-mediated ASXL1 K351 monoubiquitination machinery^29^ might achieve pathway modulation upstream.

The development of H2AK119Ub^AT^ activity-based probe represents a methodological advance for capturing transient deubiquitinase–nucleosome complexes. Previous structural studies of PR-DUB on H2AK119Ub nucleosomes have indicated that the chromatosomes with extended linker DNA and histone H1 play an important role in improving complex resolution, although PR-DUB does not interact with linker DNA or linker histone H1^35^. Our H2AK119Ub^AT^ activity-based probe, synthesized based on chemical protein semisynthesis strategies^40,41^, facilitates covalent disulfide bonding between the catalytic cysteine of PR-DUB and the isopeptide bond of the ubiquitinated histone, and enables successful cryo-EM reconstruction of BAP1/ASXL1^K351-Ub^ bound to nucleosome core particles (147-bp DNA). We anticipate that this strategy could be applied to capture and structurally elucidate other nucleosomal deubiquitinases, such as USP3-mediated deubiquitination of H2AK13/15^42–44^ and USP7-mediated deubiquitination of H3K18/23^45,46^. Notably, our previously developed H2AK15Ub^AT^ probes for RNF168 mechanism studies^47^ demonstrate the dual utility of the platform; — disulfide-trapped ubiquitinated nucleosome intermediates can co-capture E3/E2∼Ub/substrate ternary complexes, as evidenced by RNF168-mediated H2AK13/15 ubiquitination assays (**Extended Data Fig. 9**). The bidirectional trapping capability of disulfide-activated ubiquitinated nucleosome probes, which enables the capture of E2/E3 ubiquitination machinery and deubiquitinase complex, provides a unique solution to the challenge of resolving transient enzyme‒substrate interactions in chromatin ubiquitination cycles.

## Acknowledgements

L. Liu was supported by the National Key R&D Program of China (no. 2022YFC3401500), the National Natural Science Foundation of China (no. 22137005, T2488301, 92253302, 22227810), and the New Cornerstone Investigator Program. H. Ai. was supported by the Shanghai Frontiers Science Center of Drug Target Identification and Delivery (ZXWH2170101). We acknowledge the Tsinghua University Branch of China National Center for Protein Sciences (Beijing) for cryo-EM data screening and collection. We thank the Center of Protein Analysis Technology, Tsinghua University, for MS analysis.

## Author contributions

H. Ai, M. Pan and L. Liu supervised the project. H. Ai, T. Zhang, J. Zheng, Z. Tong, Z. Deng, and L. Liu proposed the idea, designed the experiments, and analyzed the results. H. Ai, Z. Tong designed the activity-based H2AK119Ub^AT^ nucleosome probe. T. Zhang, H. Ai, X. Wu, and Z. He cloned the plasmids, expressed the proteins (BAP1/ASXL1 and mutants, histones, hUba1, UBE2E1, Ub) and reconstituted the nucleosomes. T. Zhang and X. Wu prepared BAP1/ASXL1^K351Ub^. T. Zhang synthesized the H2AK119Ub histone. T. Zhang, H. Ai, and Z. Tong performed the PR-DUB (and its mutants) deubiquitination assays on H2AK119Ub nucleosomes. Z. He and Z. Deng synthesized the H2AK119Ub^AT^ nucleosome probes. H. Ai prepared the cryo-EM samples. H. Ai and Z. Tong checked the samples, and collected the cryo-EM data. H. Ai processed, determined the cryo-EM structures, and built the atomic models. T. Zhang prepared the fluorescent labelled ubiquitinated histone H2A peptide. X. Wu prepared the Ub-AMC. T. Zhang conducted the kinetic measurements. J. Zheng and T. Zhang conducted the molecular dynamic simulations. T. Zhang and X. Tian performed the tandem MS and HDX-MS analysis. T. Zhang, X. Wu, X. Tian and M. Wang analyzed the mass spectrometry data. T. Zhang, H. Ai, J. Zheng, Z. Tong, and Z. Deng collated the experimental data and prepared the figure panels and tables. Z. Deng polished the figures. H. Ai, T. Zhang, J. Zheng, Z. Deng and Z. He drafted the manuscript. H. Ai, T. Zhang, J. Zheng, Z. Tong, Z. Deng, M. Pan and L. Liu revised the manuscript. All authors read, discussed, and analyzed the manuscript.

## Figure Legends/Captions (for Extended Data Figures)

**Extended Data Figure 1.**
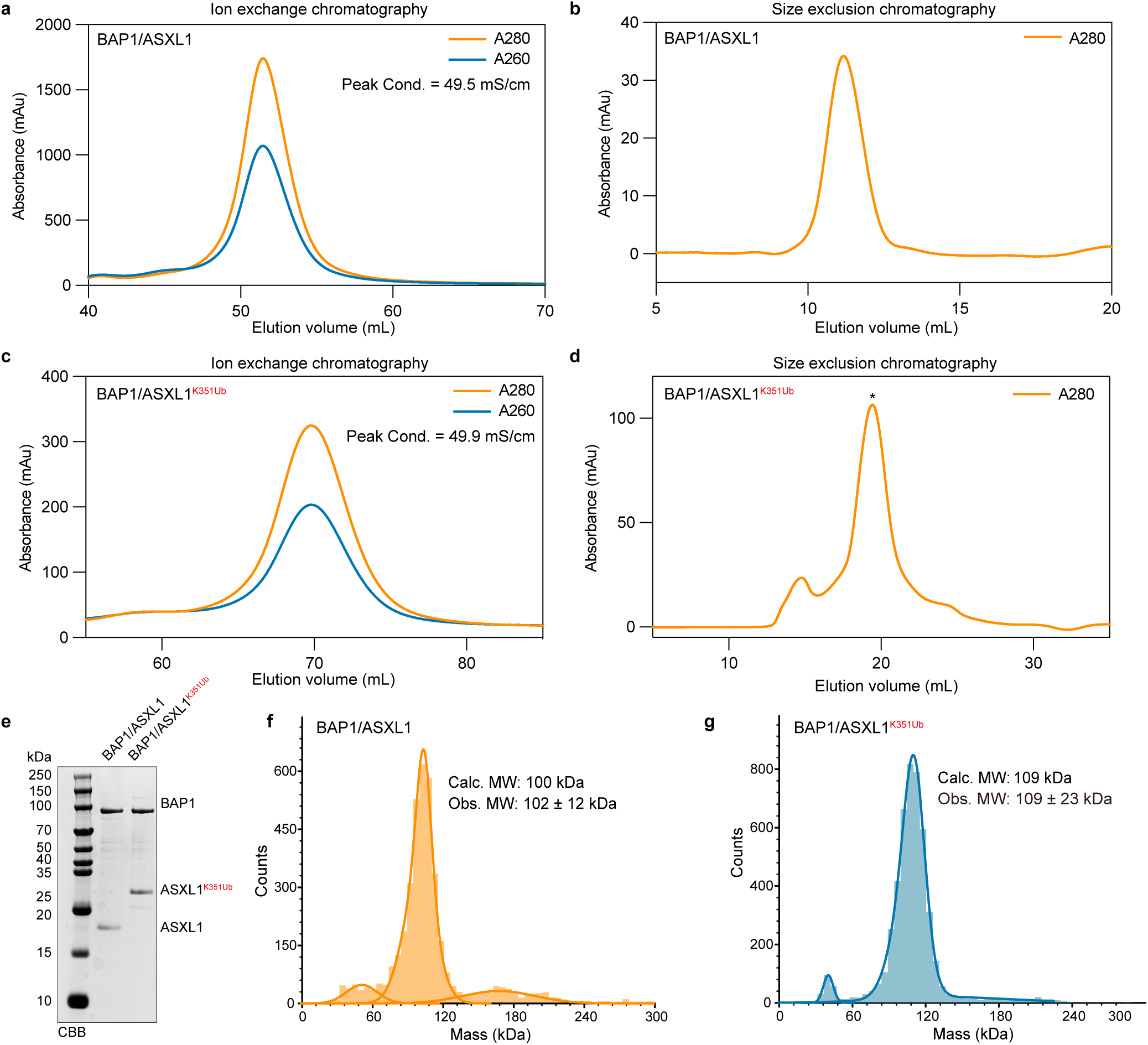
Purification and characterization of BAP1/ASXL1 and BAP1/ASXL1^K351Ub^ complexes. **a, b**, Ion exchange chromatography elution profiles for **(a)** BAP1/ASXL1 and **(b)** BAP1/ASXL1^K351Ub^. **c, d,** Size-exclusion chromatography profiles of **(c)** BAP1/ASXL1 and **(d)** BAP1/ASXL1^K351Ub^. **e,** Coomassie-stained SDS-PAGE analysis of purified BAP1/ASXL1 and BAP1/ASXL1^K351Ub^. **f, g,** Mass photometry measurements of BAP1/ASXL1 and BAP1/ASXL1^K351Ub^.

**Extended Data Figure 2.**
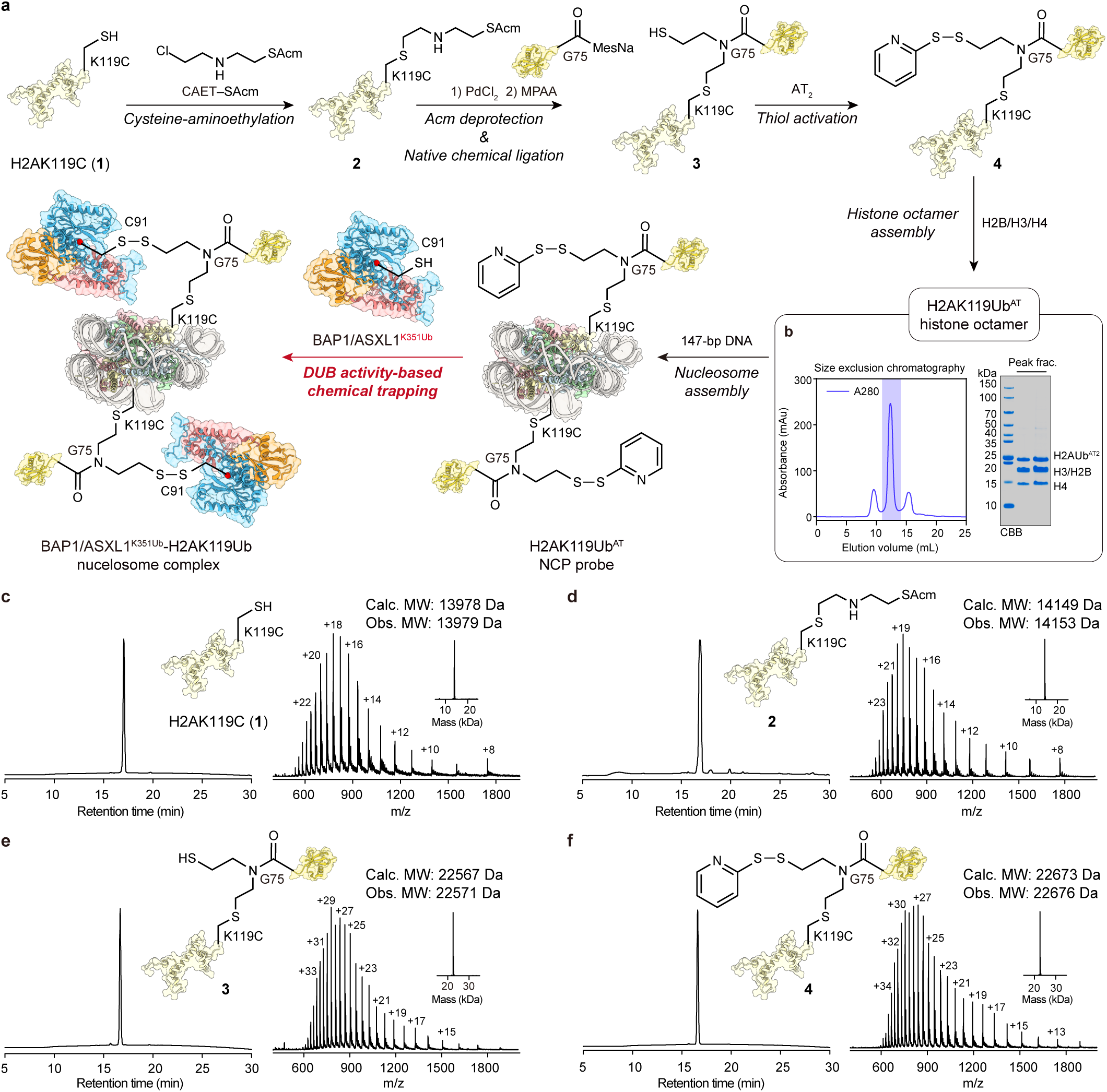
Chemical synthesis and validation of H2AK119Ub^AT^ nucleosome probe. **a**, Synthetic route of H2AK119Ub^AT^ nucleosome probe and subsequent disulfide crosslinking with BAP1/ASXL1^K351Ub^. To chemically synthesize the deubiquitinase probes, briefly, (1) the histone H2AK119C was firstly alkylated of with 2-((2-chloroethyl)amino)ethane-1-(S-acetaminomethyl)thiol (CAET), (2) removal of the ACM protecting group for subsequent native chemical ligation with Ub(1-75)MesNa thioester, and (3) activation of the thiol group via 2,2’-dithiodipyridine (AT_2_) molecules to generate histone H2AK119Ub^AT^. Then the probe was incorporated into histone octamer and nucleosome and crosslinked with BAP1/ASXL1^K351Ub^. **b,** Size-exclusion chromatography and Coomassie-stained SDS-PAGE analysis of the H2AK119Ub^AT^ octamer. **c-f,** HPLC and mass spectrometry (MS) analysis of synthetic intermediates in the synthetic route, the calculated (Calc.) molecular weight and the observation (Obs.) molecular weight were labelled.

**Extended Data Figure 3.**
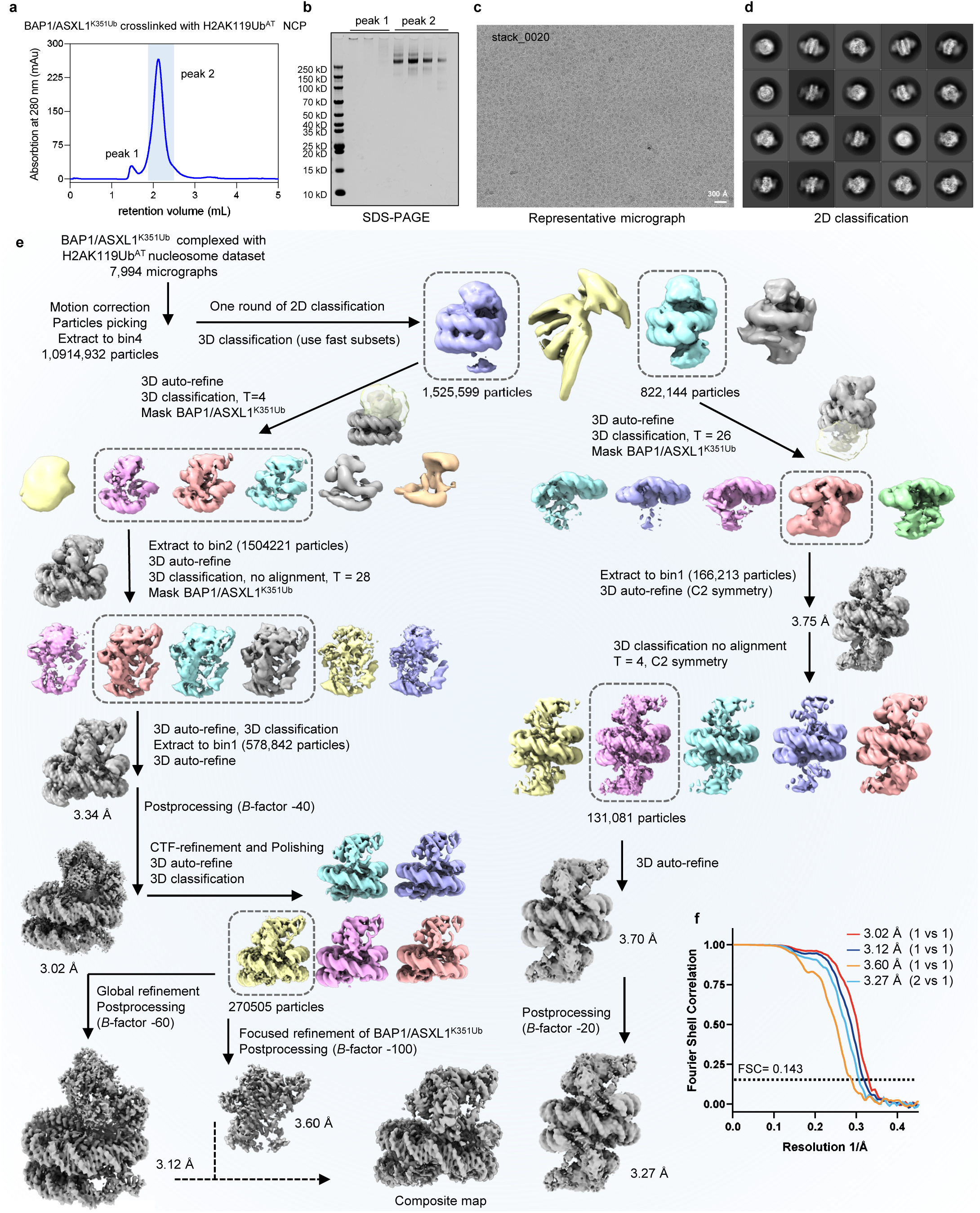
Cryo-EM data processing for the BAP1/ASXL1^K351Ub^/H2AK119Ub^AT^ nucleosome complex. **a**, Size-exclusion chromatography profile of the glutaraldehyde crosslinked BAP1/ASXL1^K351Ub^/H2AK119Ub^AT^ nucleosome complex. **b,** Coomassie-stained SDS-PAGE analysis of peak fractions from **a**. **c,** Representative cryo-EM micrograph, with all micrographs in the dataset exhibiting comparable particle density and dispersion. **d,** Representative 2D classifications. **e,** Cryo-EM data processing flowcharts. **f,** Fourier shell correlation (FSC) curves reporting global resolution for the four final maps shown in **e**.

**Extended Data Figure 4.**
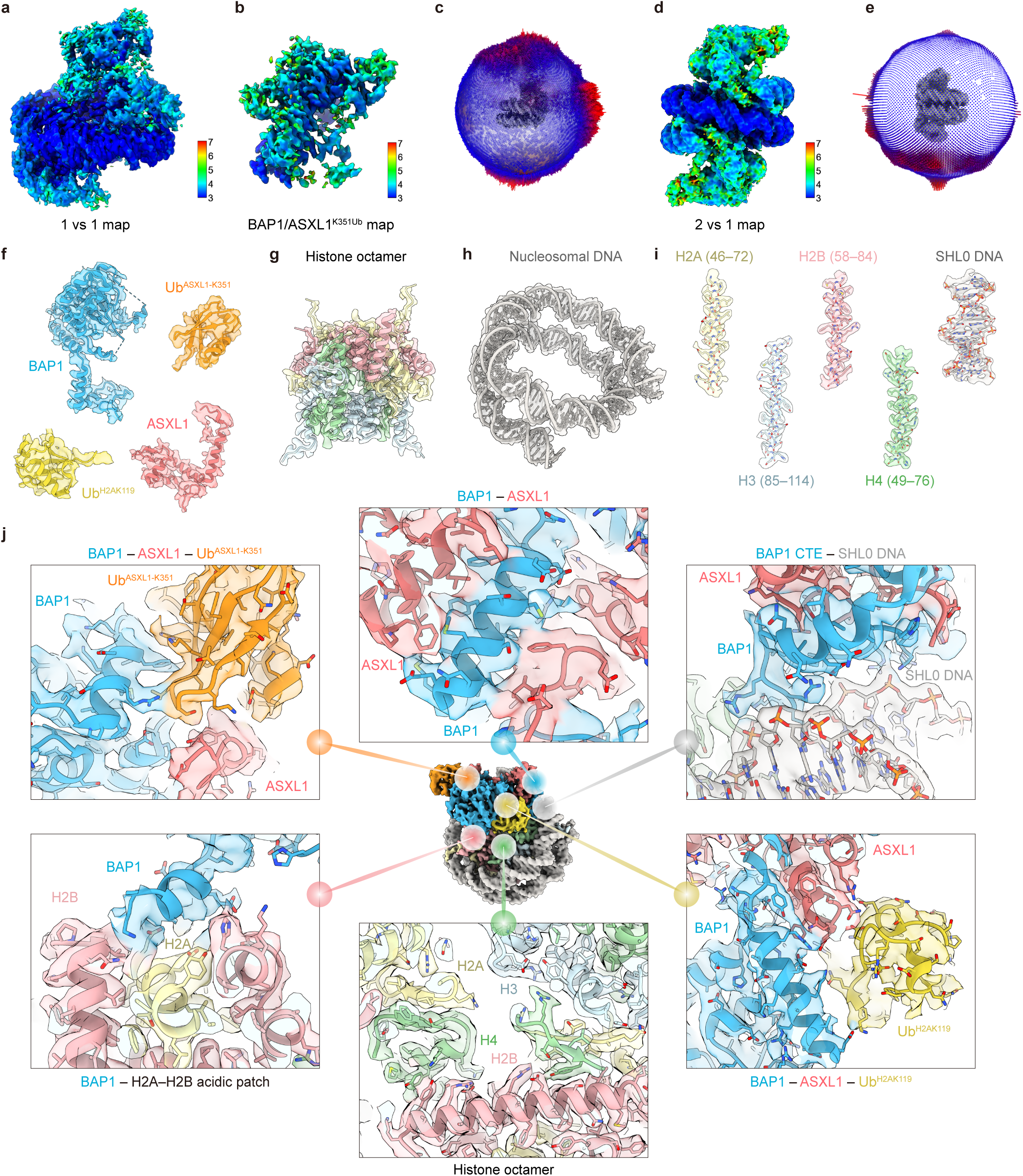
Cryo-EM map validation and interface analysis of the BAP1/ASXL1^K351Ub^/H2AK119Ub nucleosome complex. **a, b**, Local resolution maps of the **(a)** 1 vs 1 BAP1/ASXL1^K351Ub^/H2AK119Ub nucleosome complex and **(b)** BAP1/ASXL1^K351Ub^. **c, e,** Euler angle distributions of the final particles for cryo-EM reconstruction of the 1 vs 1 BAP1/ASXL1^K351Ub^/H2AK119Ub nucleosome complex conformation **(c)** and the 2 vs 1 conformation **(e)**. **d,** Local resolution map of the 2 vs 1 BAP1/ASXL1^K351Ub^/H2AK119Ub nucleosome complex map. **f-i,** Cryo-EM density superimposed with each subunit of the BAP1/ASXL1^K351Ub^/H2AK119Ub nucleosome complex. **j,** Cryo-EM density of key interfaces (Clockwise from the top): BAP1-ASXL1 heterodimer interface, BAP1 CTE domain-DNA SHL0 interface, BAP1-ASXL1-Ub^H2AK119^ interface, histone octamer, BAP1-H2A/H2B acidic patch interface, BAP1-Ub^ASXL1^ interface.

**Extended Data Figure 5.**
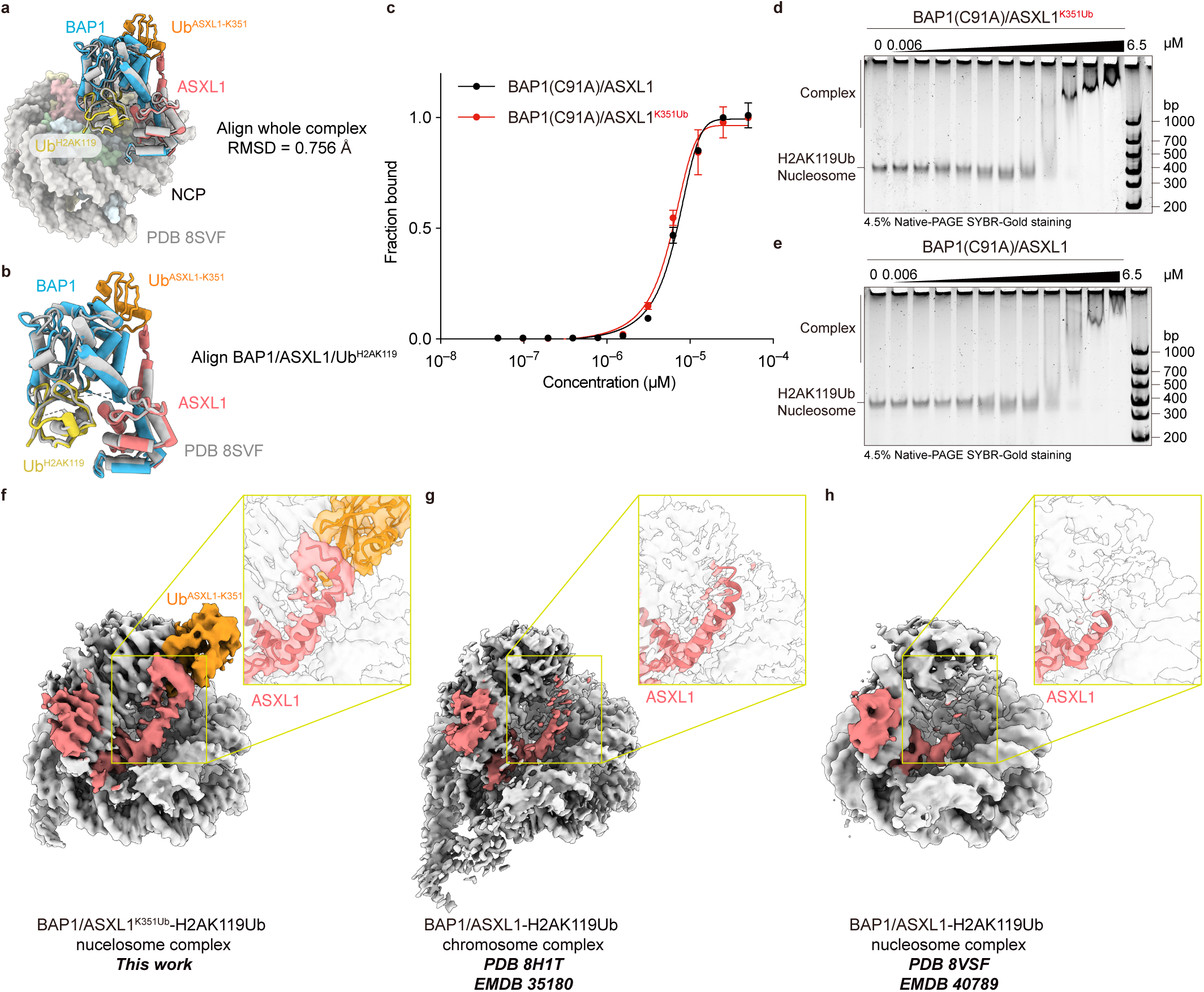
ASXL1 K351 Monoubiquitination neither alters enzyme-substrate bind mode nor strength, but stabilizes the C-terminal α-helix in the ASXL1 DEUBAD domain. a, b, Structural superposition of the BAP1/ASXL1^K351Ub^/H2AK119Ub nucleosome complex with the BAP1/ASXL1/H2AK119Ub nucleosome structure (PDB: 8VSF) a, Global alignment of the two complexes (RMSD = 0.756 Å). b, Alignment the using BAP1/ASXL1/Ub^H2AK119^ as a reference (RMSD = 1.219 Å) c, AlphaLISA analysis of catalytically inactive BAP1^C91A^/ASXL1 or BAP1^C91A^/ASXL1^K351Ub^ binding to DNA biotin-labeled H2AK119Ub nucleosomes. d, e, EMSA analysis of BAP1^C91A^/ASXL1^K351Ub^ or BAP1^C91A^/ASXL1 binding to H2AK119Ub nucleosomes. f-h, Three cryo-EM density maps of PR-DUB-H2AK119Ub nucleosome complex. The position of C-terminal α-helix of ASXL1 DEUBAD domain of each map is highlighted with a red box in each map. Corresponding EMDB entries: (f) EMD-63884 in this work, (g) EMD-35180, (h) EMD-40789.

**Extended Data Figure 6.**
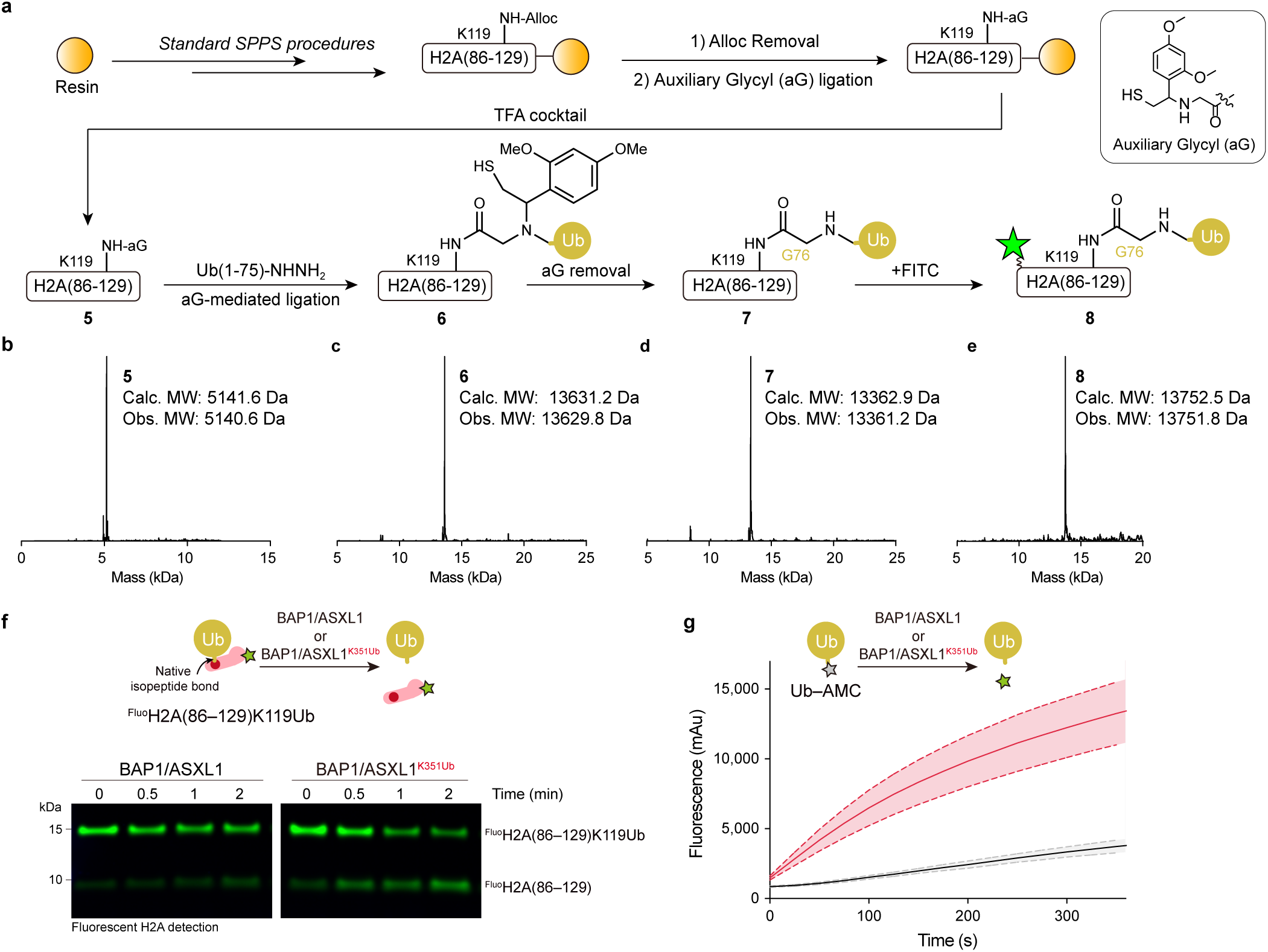
Enhanced catalytic activity of monoubiquitinated PR-DUB on non-nucleosomal substrates. **a**, Synthetic scheme for fluorescein-labeled H2A (86-129)-K119Ub peptide. **b-e,** ESI-MS characterization of synthetic intermediates, each graph confirms the molecular weight of the corresponding intermediate labelled by number. **f,** Time-course analysis reveals enhanced catalytic activity of BAP1/ASXL1^K351Ub^ compared to BAP1/ASXL1 in deubiquitinating fluorescein-labeled H2A (86-129)-K119Ub peptide, at the time point of 0, 0.5, 1 and 2 hours. **g,** Ub-AMC hydrolysis assay reveals enhanced catalytic activity of BAP1/ASXL1^K351Ub^ compared to BAP1/ASXL1. Data represent mean ± s.d. (n = 3).

**Extended Data Figure 7.**
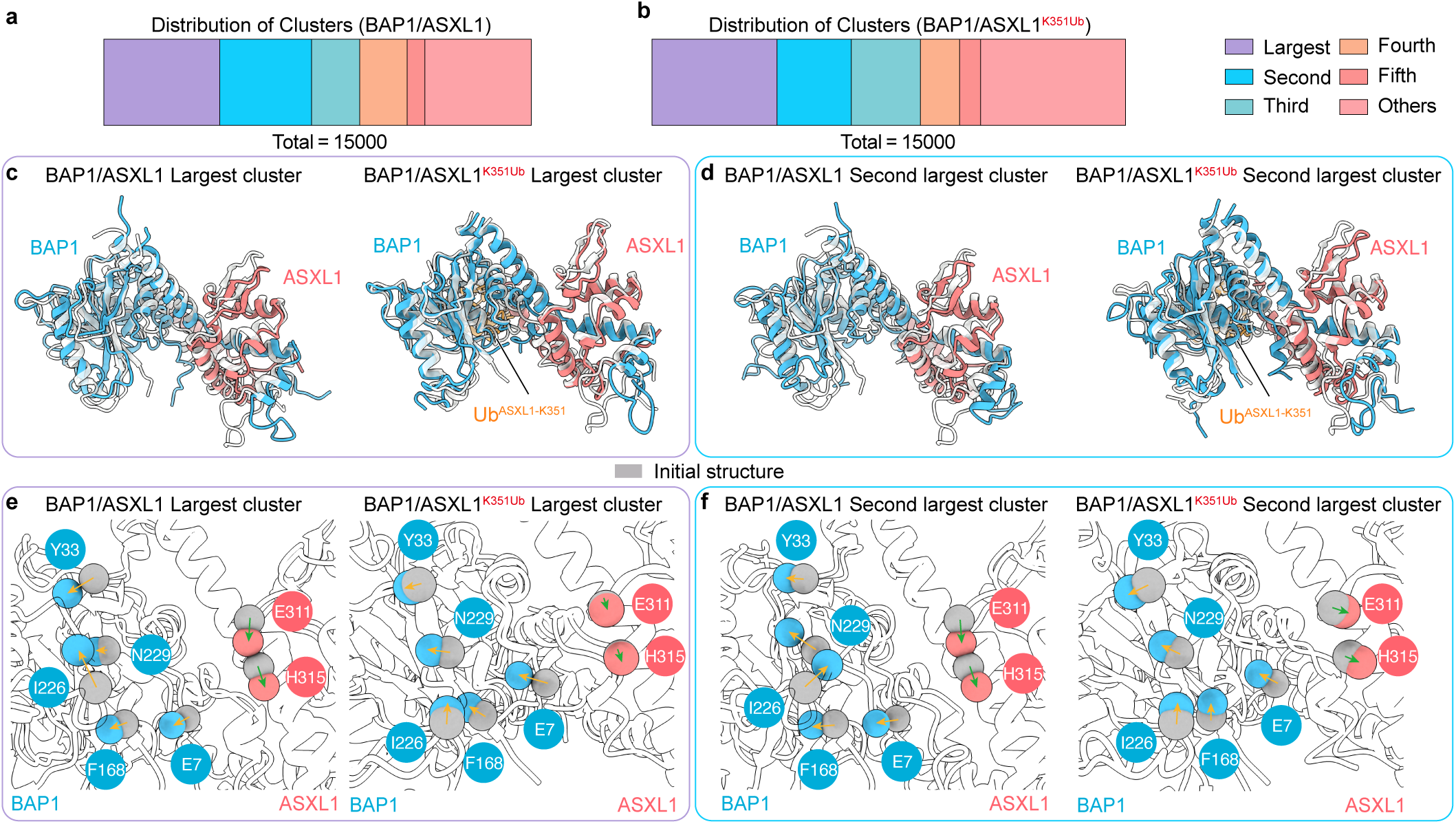
Distinct catalytic pocket patterns between BAP1/ASXL1 and BAP1/ASXL1^K351Ub^ characterized by trajectory clustering. a, b, Cluster population analysis for (a) BAP1/ASXL1 and (b) BAP1/ASXL1^K351Ub^ using centroid-based hierarchical clustering (1.5 Å cutoff). The five largest clusters and the remaining clusters (aggregated as “Other”) are color-coded as indicated separately. Total frames = 15000 per system. c–f, Structural alignment of the dominant clusters with the initial model (gray), BAP1 (cyan), ASXL1 (pink). (c) BAP1/ASXL1 and BAP1/ASXL1^K351Ub^ largest cluster; (d) Close-up views of structural repositioning in the PR-DUB catalytic pocket in largest cluster; (e) BAP1/ASXL1 and BAP1/ASXL1^K351Ub^ second largest cluster; (f) Close-up views of structural repositioning in the PR-DUB catalytic pocket in second largest cluster. Key catalytic residues (BAP1: E7, Y33, F168, I226, N229; ASXL1: E311, H315) are shown with their α-carbons shown as spheres, and gray spheres indicate initial positions to highlight conformational changes.

**Extended Data Figure 8.**
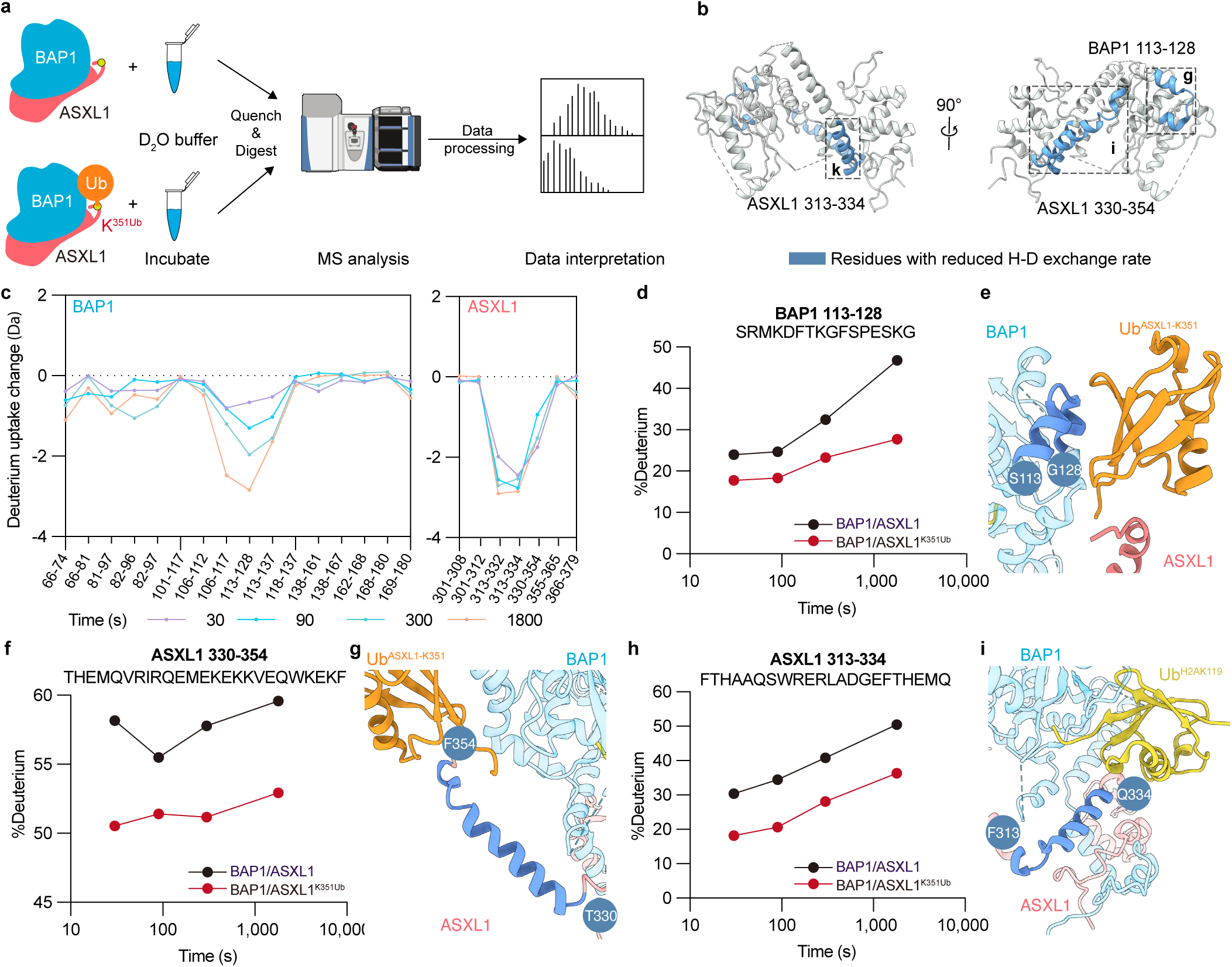
HDX-MS analysis of BAP1/ASXL1 and BAP1/ASXL1^K351Ub^. **a**, Schematic workflow of the process of HDX-MS. (Figure created with BioArt: NIAID Visual & Medical Arts. (2025/2/11). Mass Spectrometer Machine. NIAID NIH BIOART Source. bioart.niaid.nih.gov/bioart/579). **b,** Structural mapping of peptides exhibiting reduced deuterium uptake (blue) in BAP1/ASXL1. Two perspectives were shown. **c,** Deuterium uptake profiles for peptides with significant difference. The full dataset is in **Supplementary** Fig. 2. **d-i,** Detailed analysis of deuterium uptake rate reduced peptides. **d, f, h,** Time-resolved deuterium uptake profile of **(d)** BAP1 residues 113-128, **(f)** ASXL1 residues 330-354, **(h)** ASXL1 residues 313-334. **e, g, i,** Peptides with reduced deuterium uptake (blue) mapped onto our determined BAP1/ASXL1^K351Ub^-H2AK119Ub nucleosome structure (PDB: 9U5U). **e,** BAP1 residues 113-128 and surround regions. **g,** ASXL1 residues 330-354 and surround regions. **i,** ASXL1 residues 313-334 and surround regions.

**Extended Data Figure 9.**
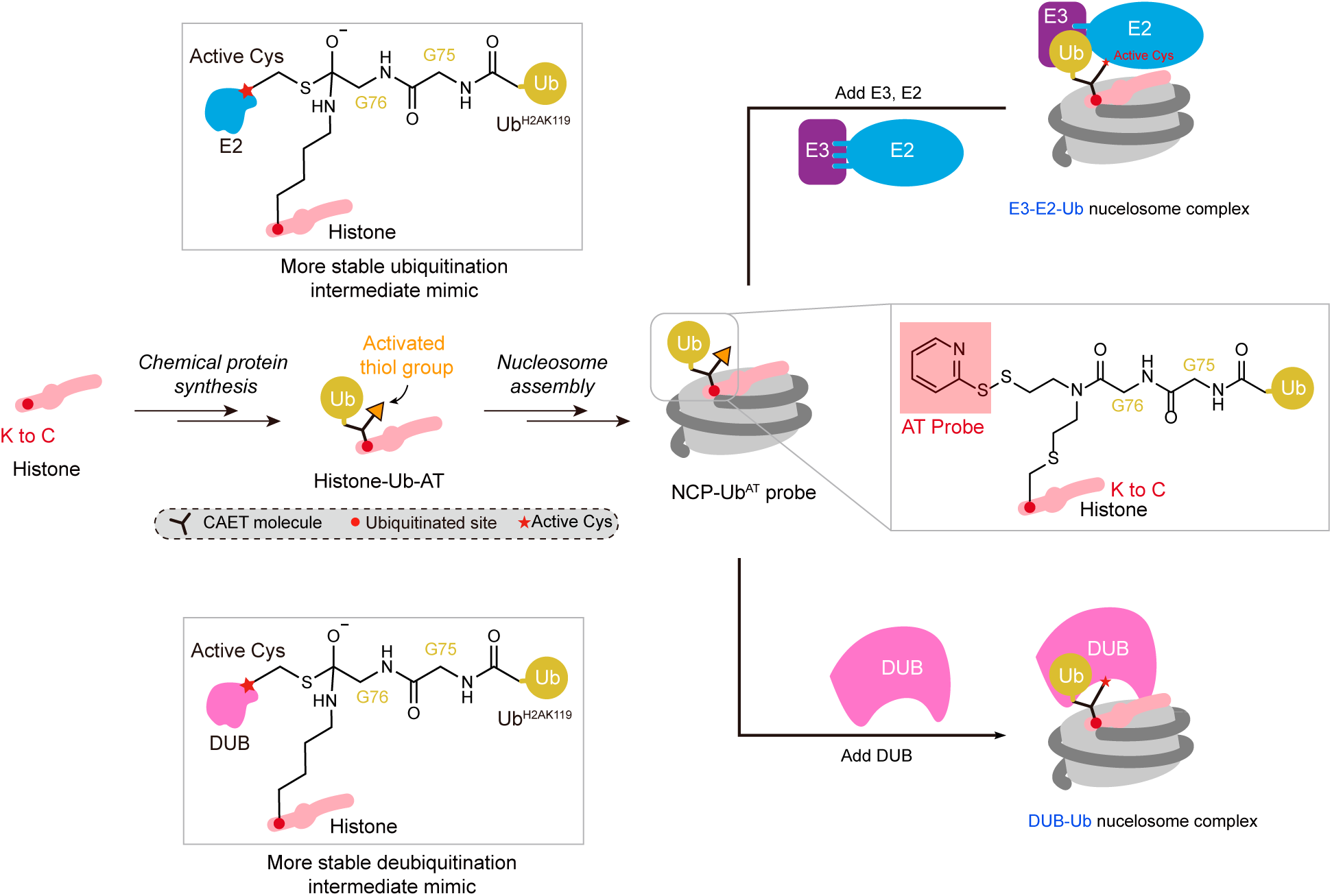
Functional versatility of the disulfide-trapped ubiquitinated nucleosome intermediate probe. Schematic illustrating the bifunctional utility of the H2AK119UbAT probe for trapping E2-E3 ubiquitin ligase complexes during ubiquitin transfer and deubiquitinases (DUBs) during substrate hydrolysis. Structural models of the probe mimicking ubiquitination (E2-E3 trapping) and deubiquitination (DUB trapping) intermediates are shown. A lysine-to-cysteine histone mutant is conjugated to ubiquitin via a CAET molecule, which introduces an activated thiol group for trapping. For the trapping of E2-E3 complex, the nucleosome probe is incubated with E2-E3 complex, and crosslinks with the active cysteine of E2 enzyme through a disulfide bond. For the trapping of DUB, the probe crosslinks with the catalytic cysteine of DUB.

**Supplementary Figure 1.**
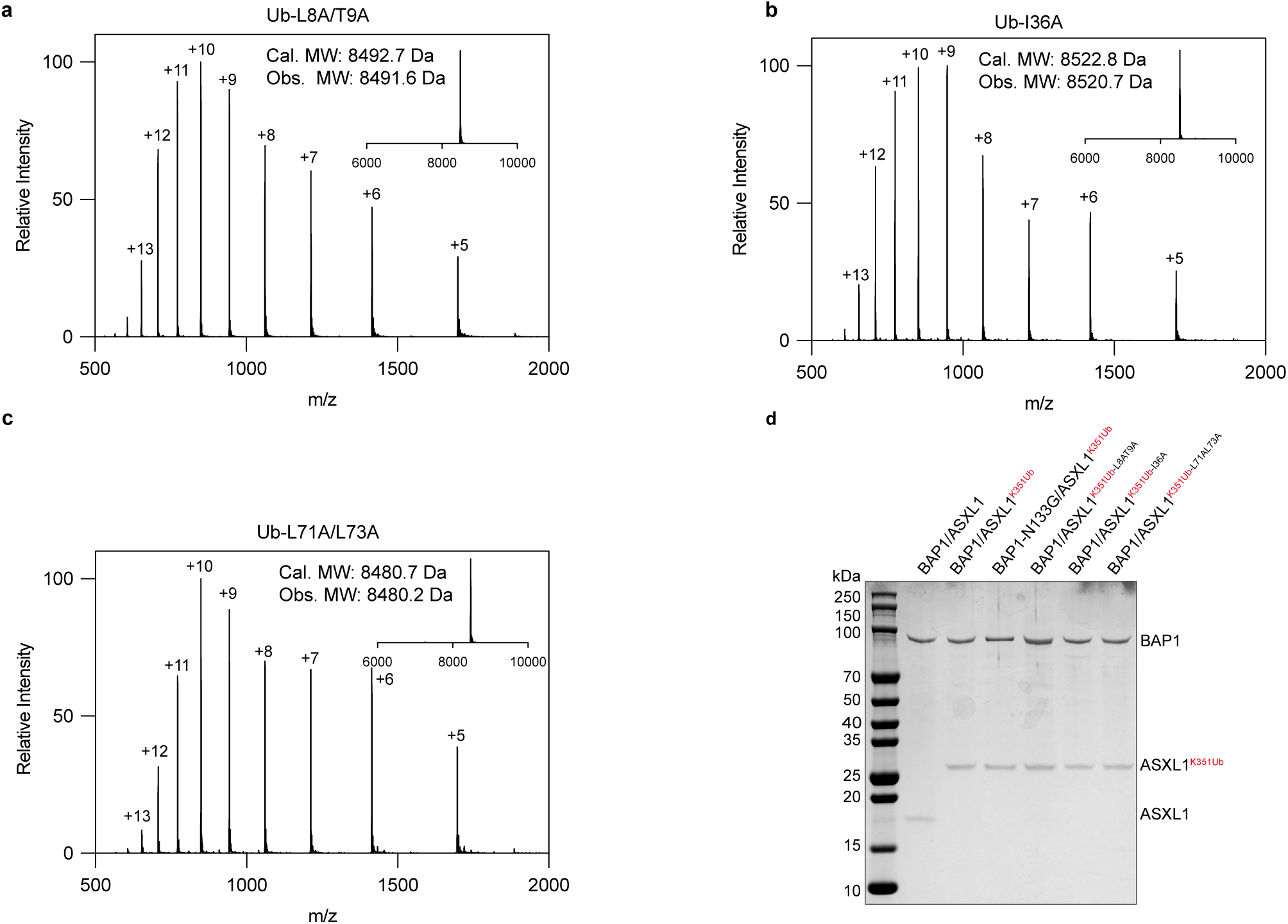
Validation of ubiquitin and BAP1/ASXL1K351Ub mutants. a-c, Mass spectrometry confirmation of ubiquitin mutant molecular weights: (a) Ub-L8A/T9A, (b) Ub-I36A, (c) Ub-L71A/L73A. d, Coomassie-stained SDS-PAGE analysis of wild-type (WT) BAP1/ASXL1, WT-BAP1/ASXL1K351Ub and its mutants used in in vitro deubiquitination assays.

**Supplementary Figure 2.**
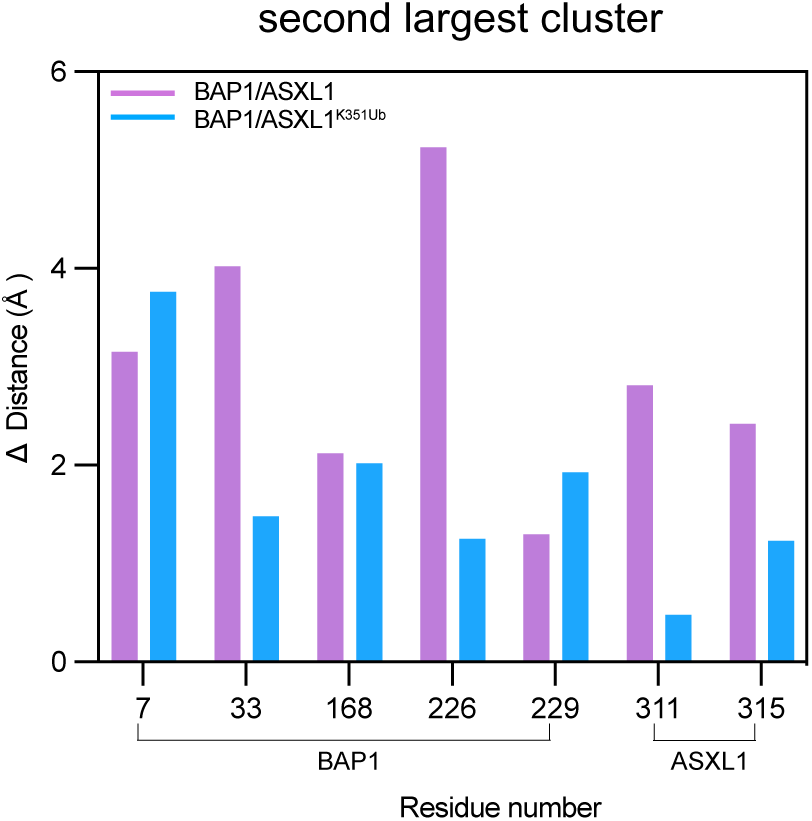
**a**, α-carbon distance comparison for between the initial structure and the second most populated MD cluster structure for the seven key residues.

**Supplementary Figure 3.**
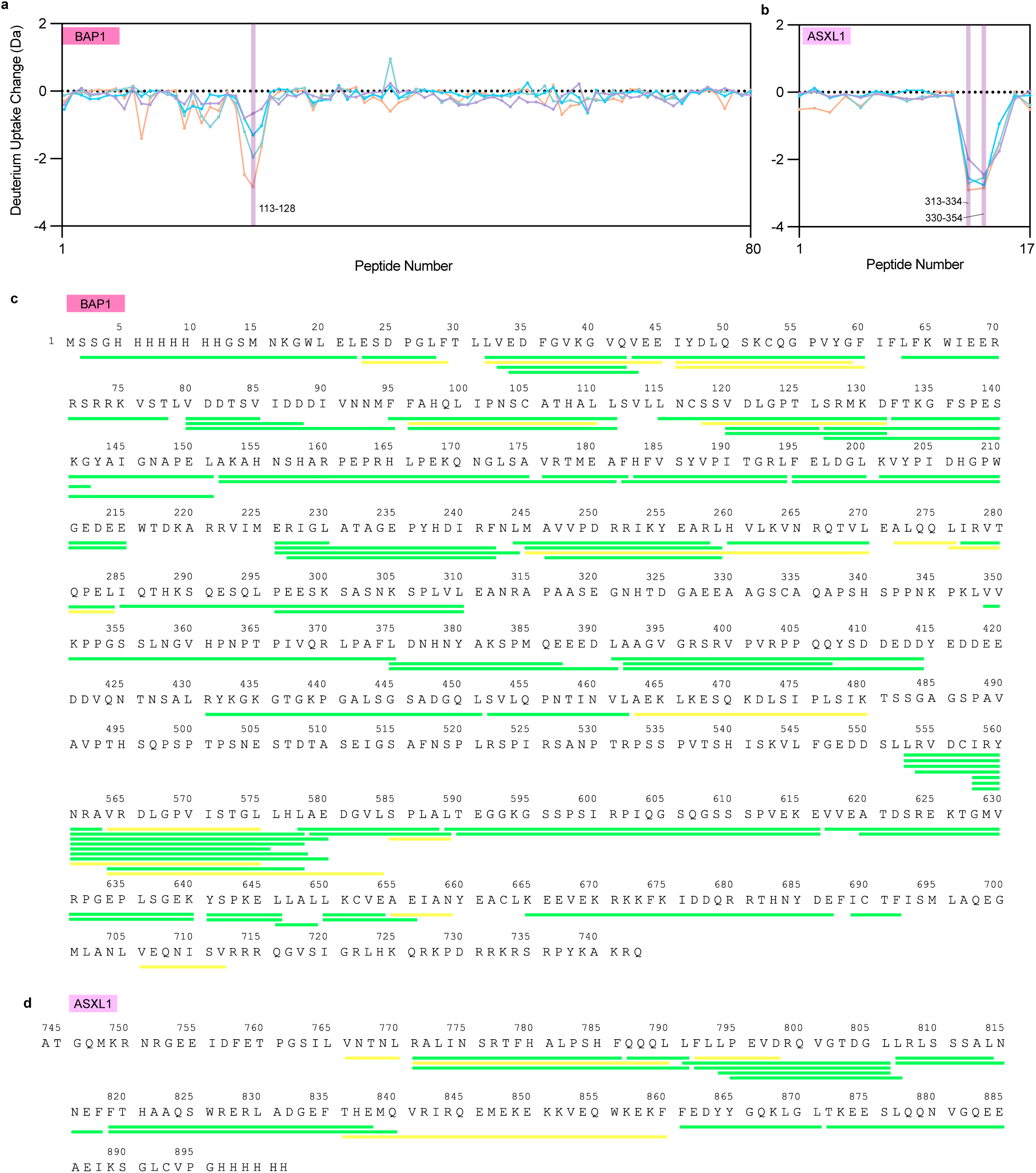
Deuterium uptake change and coverage maps form PR-DUB HDX experiments. a, b, Deuterium uptake change for (a) BAP1 and (b) ASXL1. c, d, HDX coverage maps for (c) BAP1 and (d) ASXL1. The His-tag sequence was included in the HDX-MS analysis. ASXL1 residue numbering starts at 745 due to sequential concatenation with BAP1 during data processing. Peptides with high confidence (green) and medium confidence (yellow) are shown.

**Extended Data Table 1.** Cryo-EM data collection, refinement, and validation statistics.

|  | BAP1/ASXL1<sup>K351Ub</sup> in complex with<br>H2AK119Ub<sup>AT</sup> nucleosome (1:1)<br>(EMD-63884, PDB: 9U5U) | BAP1/ASXL1<sup>K351Ub</sup> in complex with<br>H2AK119Ub<sup>AT</sup> nucleosome (2:1)<br>(EMD-63885) |
| --- | --- | --- |
| Data collection and processing |  |  |
| Magnification | 64,000 |  |
| Voltage (kV) | 300 |  |
| Spherical aberration (mm) | 2.7 |  |
| Detector | K3 |  |
| Electron exposure (e<sup>-</sup>/Å<sup>2</sup>) | 50.0 e<sup>-</sup>, 32 frames |  |
| Defocus range (μm) | -1.0 to -2.0 |  |
| Pixel size (Å) | 1.0979 |  |
| Box size (pixel) | 256 |  |
| Micrographs (no.) | 47,994 |  |
| Initial particle images (no.) | 1,0914,932 |  |
| Symmetry imposed | C1 | C2 |
| Final particle images (no.) | 270,505 | 131,081 |
| Map resolution (Å) | 3.12 Å | 3.27 Å |
| FSC threshold | 0.143 | 0.143 |
| Refinement |  |  |
| Initial model use (PDB code) | 1UBQ, 7XD1, 8H1T |  |
| Model resolution range (Å) | 3.0-8.0 |  |
| Map sharpening B factor (Å<sup>2</sup>) | -60 |  |
| Model composition |  |  |
| Nonhydrogen atoms | 16,392 |  |
| Protein residues | 1,295 |  |
| Nucleotides | 294 |  |
| B factors (Å<sup>2</sup>) |  |  |
| Protein | 16.65/322.91/82.84 |  |
| Nucleotide | 34.95/266.97/105.04 |  |
| R.m.s. deviations |  |  |
| Bond lengths (Å) | 0.009 |  |
| Bond angles (°) | 0.646 |  |
| Validation |  |  |
| MolProbity score | 1.79 |  |
| Clashscore | 7.98 |  |
| Poor rotamers (%) | 1.72 |  |
| Ramachandran plot |  |  |
| Favored (%) | 97.00 |  |
| Allowed (%) | 3.00 |  |
| Disallowed (%) | 0.00 |  |

**Supplementary Table S1.**
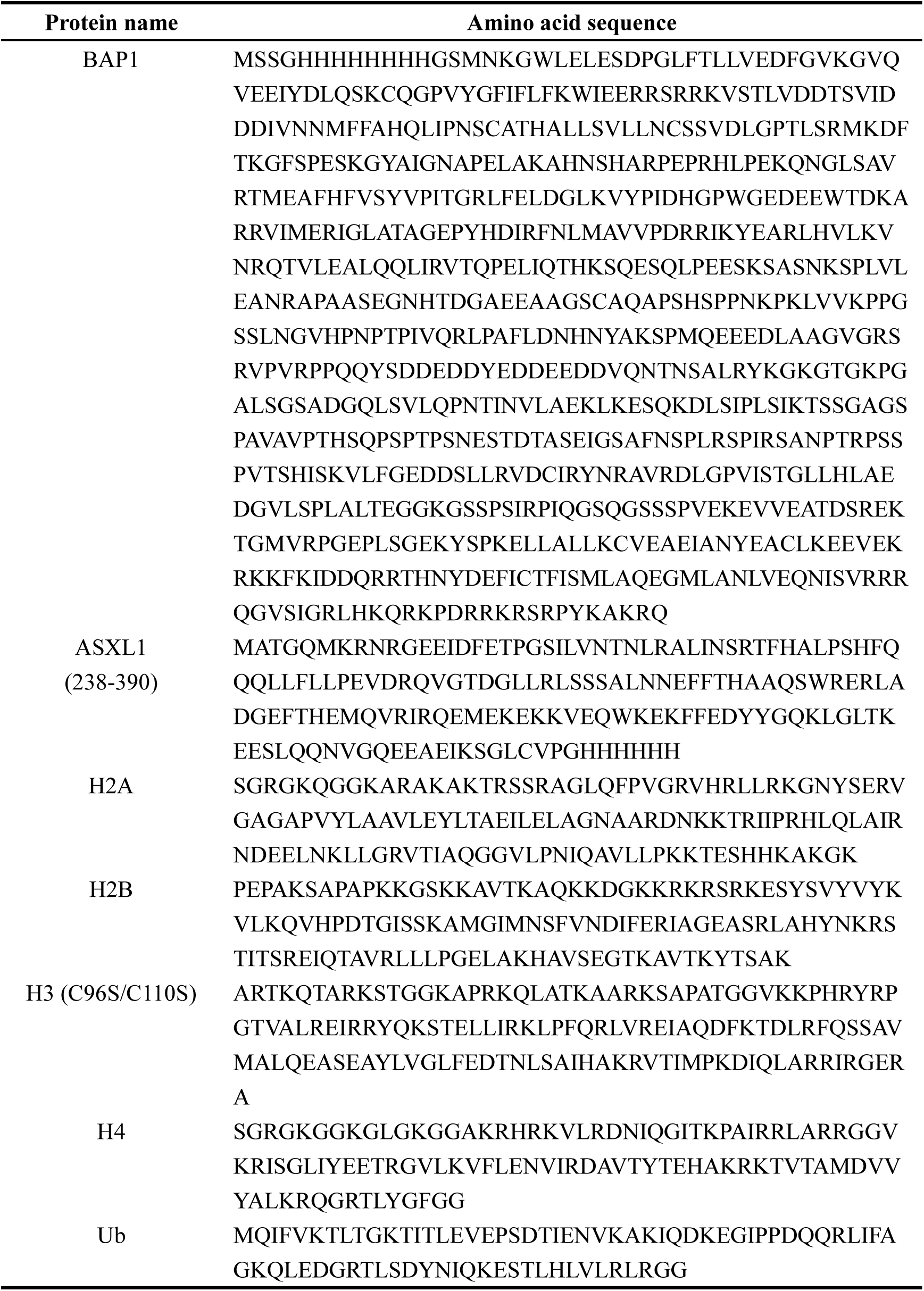
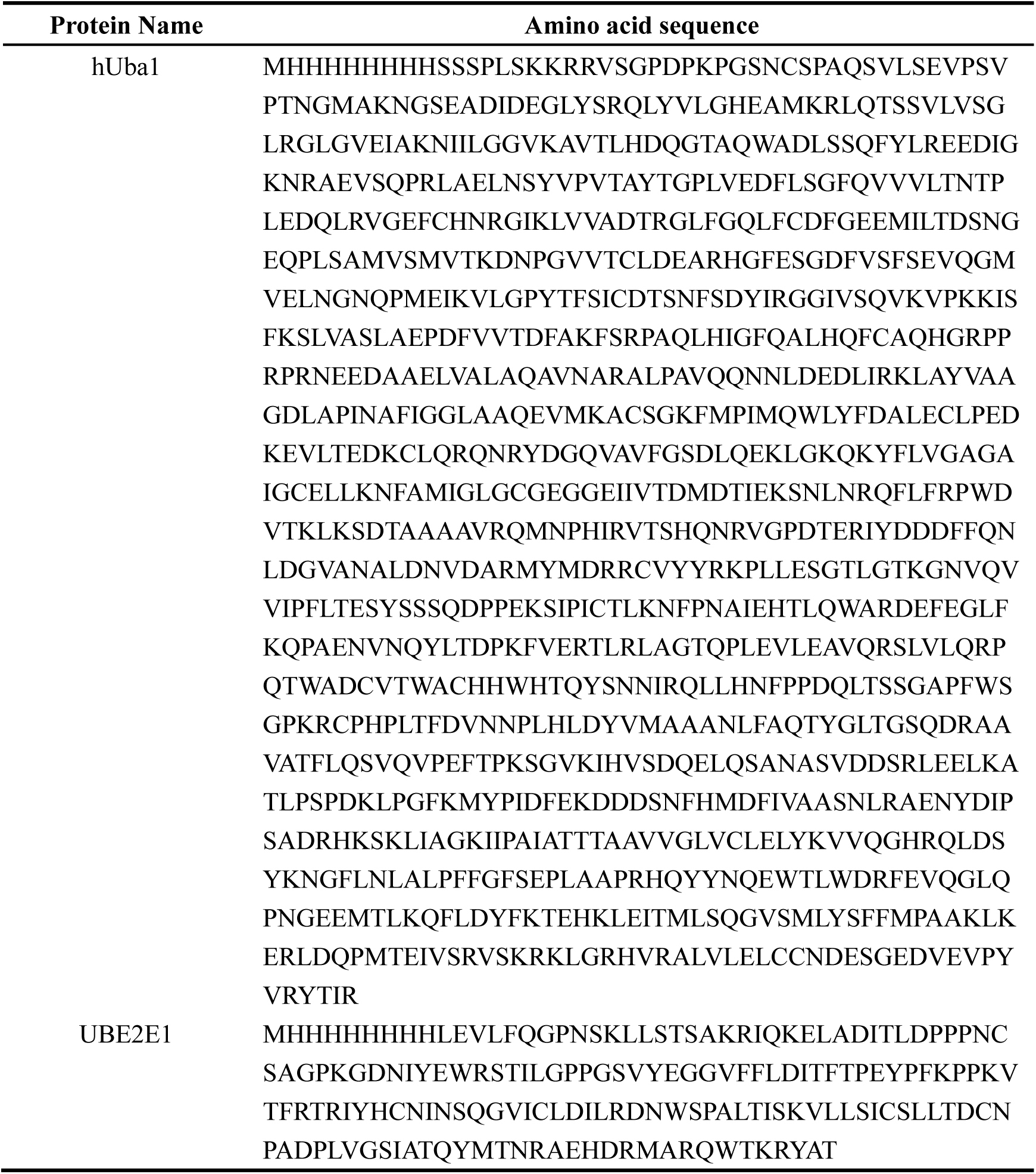
Protein sequences.

## Methods

### Plasmid construction

The human BAP1 gene (residues 1-729) and the human ASXL1 gene (residues 238-390) were synthesized by GenScript (Nanjing, China), with sequences optimized for expression in *Escherichia coli*. The BAP1 gene was cloned into a modified pET28a expression vector containing an N-terminal 6xHis–SUMO tag, while the ASXL1 gene was inserted into the pET-22b vector with a C-terminal 6×His tag. For the BAP1 mutants, the N-terminal 8×His tag was replaced to reduce purification steps. All the mutations were introduced by standard site-directed PCR mutagenesis.

The genes encoding histones (H2A, H2B, H3 (C96S/C110S), H4) and ubiquitin were cloned into the pET-22b vector without any affinity tag. The human Uba1 gene was cloned into the pET-3a vector with an N-terminal 6×His tag. The genes encoding human UBE2E1 were constructed as previously reported^34^.

### Protein expression and purification

BAP1/ASXL1 were co-expressed in *E*. coli and purified using Ni-NTA affinity chromatography (LABLEAD Inc., Cat: N30210) followed by ion exchange chromatography (HiTrap Heparin column, GE Healthcare) and size-exclusion chromatography (Superose 6 Increase 10/300 GL column, GE Healthcare). First, BAP1 and ASXL1 plasmids were co-transformed into *E*. coli BL21(DE3) competent cell (Transgen, Cat: CD601-02), and cultured at 37℃ in Luria Broth (LB) medium (Coolaber, Cat: PM0010-5kg) containing 50 μg/mL kanamycin and 50 μg/mL ampicillin. After the OD600 reached 0.6, the cell culture was induced with 0.4 mM isopropyl β-D-thiogalactopyranoside (IPTG; LABLEAD Inc., Cat: 0417) for 16 hours at 16°C. Cells culture was harvest by centrifugation at 5000 ×g for 30 minutes at 4℃. The cell pellet was resuspended by washing buffer (50 mM HEPES, 1 M NaCl, 20 mM imidazole, 1 mM DTT, 20% glycerol, pH 7.5) and then lysed by sonication. Lysates were centrifuged at 20,000 ×g for 30 minutes at 4℃. The supernatant was loaded onto Ni-NTA that had been pre-equilibrated with wash buffer, incubated for 2 h at 4℃. Then Ni-NTA beads washed by 10 column volumes of washing buffer. Subsequently, the protein was eluted by elution buffer (50 mM HEPES, 1 M NaCl, 300 mM imidazole, 1 mM DTT, 20% glycerol, pH 7.5). The eluted proteins were dialyzed against the ion-exchange (IEX) Buffer A (50 mM HEPES, 50 mM NaCl, 1 mM DTT, 5% glycerol, pH 7.5) overnight at 4°C. His6-SUMO tags on BAP1 were cleaved by adding home-made Ulp1 protease. The proteins were subsequently loaded into a 5 mL heparin affinity column (HiTrap Heparin HP, Cytiva) equilibrated with the IEX Buffer A then eluted by a gradient of the IEX Buffer A and IEX Buffer B (50 mM HEPES, 1 M NaCl, 1 mM DTT, 5% glycerol, pH 7.5). The peak fractions analyzed by 4-12% SDS–PAGE (SurePAGE, GenScript, Cat: M00654) were mixed and concentrated and further purified by size-exclusion chromatography (Superdex 6 Increase 10/300, Cytiva) pre-equilibrated by SEC buffer (20 mM HEPES, pH 7.5, 300mM NaCl, 1 mM DTT, 5% glycerol). The peak fractions were analyzed by SDS-PAGE and target fractions were rapidly frozen and stored at −80 ℃. For the BAP/AXSL1 mutants, the protein was eluted from Ni-NTA and directly purified by affinity chromatography (HiTrap Heparin HP, Cytiva) and size-exclusion (Superdex 6 10/300 GL column, Cytiva) chromatography.

The histones (H2A, H2B, H3, H4) and Ub or its mutants were expressed and purified as described in previous studies^48^. The histone H2A with a native isopeptide bond linked to Ub at its K119 site was chemically synthesized as previously described^49^. Human Uba1 and human UBE2E1 were expressed in *E*. coli and purified as previously reported^34^

All the protein sequences were provided in the Supplementary Table S1.

### Preparation of BAP1/ASXL1^K351Ub^ by UBE2E1 strategy

1 μM hUba1, 5 μM UBE2E1, 10 μM freshly purified BAP1/ASXL1 or mutants, and 15 μM Ub were incubated in the reaction buffer (50 mM HEPES, 150 mM NaCl, 10 mM ATP, 5 mM MgCl_2_, pH 7.5) at 37 ℃. The reaction process was monitored using Coomassie-stained SDS-PAGE. Following the complete ubiquitination of ASXL1 subunits, the reaction mixture was subjected to heparin affinity column (HiTrap Heparin HP 5/50 GL, Cytiva) with equilibrated IEX buffer A (50 mM HEPES, 50 mM NaCl, 1 mM DTT, 5% glycerol, pH 7.5), then eluted by a gradient of the IEX Buffer A and IEX Buffer B (50 mM HEPES, 1 M NaCl, 1 mM DTT, 5% glycerol, pH 7.5). Target fractions of ubiquitinated BAP1/ASXL1 (BAP1/ASXL1^K351Ub^) were collected and rapidly frozen for storage at −80 ℃.

### Western Blot

SDS-PAGE analysis was performed and proteins were transferred to a PVDF membrane using the eBlot protein transfer system (eBlot L1, GenScript). The membrane was then incubated with blocking buffer (Beyotime, Cat: P0252) for 1 h at room temperature, followed by incubating with a 1:5000 diluted anti-His tag antibody (Immunoway, Cat: YM3004). HRP* goat anti mouse IgG (H+L) (Immunoway, Cat: S0001) was 1:10000 diluted and used as the secondary antibody. ECL reagent (Bio-Rad, Cat: 1705060) was dropwise added for chemiluminescence imaging using the GelDoc Go Imaging System (Bio-Rad).

### Tandem Mass Spectrometry

The Coomassie-stained gel band corresponding to ASXL1^K351Ub^ was excised after SDS-PAGE analysis and then underwent to in-gel trypsin digestion overnight at 37 ℃. The digested peptides were extracted using a solution containing 0.1% trifluoroacetic acid and 50% aqueous acetonitrile, followed by drying under vacuum centrifugation (SpeedVac, Thermo Scientific). LC-MS/MS analysis was performed on an Ultimate nanoflow HPLC system (Dionex) coupled to an LTQ Orbitrap Velos mass spectrometer (Thermo Scientific). A gradient analysis for 45 min was performed to separate the peptides. Data-dependent acquisition was performed with Xcalibur 3.0 software. A single full MS scan (350-1550 m/z, 120,000 resolution) was acquired, followed by top-speed MS/MS scans in the Orbitrap. Raw data was processed using Proteome Discoverer (Version PD1.4, Thermo Fisher) based on the target sequence. The search parameters were as follows: peptide mass tolerance of 20 ppm, MS/MS tolerance of 0.8 Da.

### Mass Photometry

The mass photometry (MP) measurements were carried out on a Two^MP^ mass photometer (Refeyn) at 25 ℃. Proteins were diluted to 10 nM in MP buffer (50 mM HEPES, 150 mM NaCl, 1 mM DTT, pH 7.5). The standard mass curve was calibrated with beta amylase (Sigma-Aldrich, Cat: A8781). Movies were recorded for 60 s (30,000 frames) using AcquireMP software and default settings. DiscoverMP software was used to analysis the data.

### Reconstitution of histone octamers and nucleosomes

To reconstitute histone octamers, four kinds of histone dry powders (equivalent ratio, H2AK119Ub: H2B: H3 (C96S/C110S): H4 at 1.05:1.05:1:1) were weighed and dissolved in denaturation buffer (6 M guanidine hydrochloride, 20 mM Tris, pH 7.5). Histone solutions were then mixed and transferred to a dialysis tube (D-Tube Dialyzer Maxi, Milipore, Cat:71508), against refolding buffer (10 mM Tris, 2 M NaCl, 1 mM EDTA, pH 7.5). The dialysate was changed once every 6∼8 h. The fully assembled histone octamers was purified by size-exclusion chromatography (Superdex 200 Increase, 10/300 GL, Cytiva).

To reconstitute nucleosome, purified octamer was mixed with 147bp 601-Widom DNA (1:1 molar ratio) in in the refolding buffer. The mixture was transferred to a dialysis tube (D-Tube Dialyzer Maxi, Milipore, Cat:71508) against a refolding buffer. The HE buffer (10 mM HEPES, 1 mM EDTA, pH 8.0) was gradually introduced to the dialysate using a peristaltic pump (∼0.5 mL/min) to lower the salt concentration to 0.2∼0.4 M. The mixture was subsequently dialyzed into the HE buffer for 4 h. The nucleosome was further purified by anion exchange chromatography (DEAE column, Tosoh TSKgel) with a gradient of the buffer A (10 mM HEPES, 1 mM EDTA, pH 7.5) and buffer B (10 mM HEPES, 1 mM EDTA, 1 M NaCl, pH 7.5).

### In vitro deubiquitination assay

For the time-resolved nucleosomal deubiquitination experiments, 100 nM BAP1/ASXL1 or BAP1/ASXL1^K351Ub^ complex or their mutants was incubated with 800 nM H2AK119Ub nucleosome in a DUB reaction buffer (50 mM HEPES, 100 mM NaCl, 2 mM MgCl_2_, 1 mM DTT, pH 7.5) at 30℃ and aliquots were taken at 0, 0.5, 1, 2, and 5 min and quenched by adding 4×LDS buffer containing 400 mM DTT., The samples were resolved by 4-12% NuPAGE Bis-Tris (Thermo Fisher Scientific, Cat: NP0336BOX) gels and stained by Coomassie Brilliant blue. The reaction conversion rate was estimated through densitometric analysis of Coomassie-stained H2AUb and H4 bands by Image Lab-6.0.1 (Bio-Rad software), and calculated as the H2AUb/H4 ratio at specific times divided by the initial H2AUb/H4 ratio. The conversion-time curves for deubiquitination were plotted using Prism 10 (GraphPad software).

For the concentration gradient nucleosomal deubiquitination experiments, serially diluted BAP1/ASXL1 or BAP1/ASXL1^K351Ub^ complex (100 nM, 50 nM, 25 nM, 12.5 nM, and 6.25 nM) was incubated with 800 nM H2AK119Ub nucleosome in the DUB reaction buffer for 5 min at 30 ℃. Reactions were quenched by 4×LDS buffer (NuPAGE™, Invitrogen) containing 400 mM DTT and reaction conversion rate were analyzed as described above.

For the fluorescein-labeled H2A (86-129)-K119Ub peptide (FITC-H2A-Ub) deubiquitination assay, 100 nM BAP1/ASXL1 or BAP1/ASXL1^K351Ub^ was incubated with 10 μM FITC-H2A-Ub and aliquots were taken at 0, 0.5, 1, 2 min at 30 ℃. Reactions were quenched by 4×LDS buffer containing 400 mM DTT and reaction conversion rate were analyzed as described above. For Michaelis-Menten kinetics, 50 nM BAP1/ASXL1 or BAP1/ASXL1^K351Ub^ was incubated with serially diluted FITC-H2A-Ub (40 μM, 20 μM, 10 μM, 5 μM, 2.50 μM, 1.25 μM). Initial velocities were monitored by SDS-PAGE analysis, and plotted against substrate concentration and fitted to the Michaelis-Menten model in Prism 10 (GraphPad software).

### Ub-AMC hydrolysis assay

20 μL of 25 nM BAP1/ASXL1 or BAP1/ASXL1^K351Ub^ and 20 μL of 20 μM Ub-AMC in DUB buffer (50 mM HEPES, 100 mM NaCl, 2 mM MgCl_2_, 1 mM DTT, pH 7.5) were mixed and incubated in white 384-well plates (OptiPlate, Revvity) at a 40 μL final volume. Fluorescence (λ_ex_ = 360 nm, λ_em_ = 460 nm) was monitored every 25 s for 10 min using a Synergy H1 plate reader (BioTek).

For Michaelis-Menten kinetics, 20 μL of 50 nM BAP1/ASXL1 or BAP1/ASXL1^K351Ub^ and Ub-AMC with varied concentration (80 μM, 40 μM, 20 μM, 10 μM, 5 μM, 2.50 μM) were mixed and incubated in white 384-well plates (OptiPlate, Revvity). Fluorescence (λ_ex_ = 360 nm, λ_em_ = 460 nm) was monitored every 25 s for 10 min using a Synergy H1 plate reader (BioTek). Initial velocities were derived from fluorescence measurements using an AMC standard curve. Initial reaction rates were plotted against substrate concentration and fitted to the Michaelis-Menten model via non-linear regression in Prism 10 (GraphPad software).

### Chemical synthesis of H2AK119Ub^AT^ probe

Lyophilized H2AK119C peptide was solubilized in reaction buffer (100 mM HEPES, 6 M Gn·HCl, pH 7.8) at a final concentration of 1 mM. Then tris(2-carboxyethyl) phosphine (TCEP) was introduced was added to reach a final concentration of 5 mg/mL, followed by ultrasonic dissolution. Subsequently, a 30-fold molar excess of CAET-Acm molecules (2-((2-chloroethyl) amino) ethane-1-(S-acetaminomethyl)thiol)^50,51^ was added, and the pH was adjusted to 8.5. The reaction solution was stirred at 30 °C for 16 hours. H2AK119C-CAET-SAcm was isolated through semi-preparative RP-HPLC (XB-C18 column, Welch) purification and lyophilized to yield the powdered form.

Lyophilized H2AK119C-CAET-SACm powder was dissolved in ligation buffer (100 mM HEPES, 6 M Gn·HCl, pH 7.8) at a final concentration of 1 mM. 5 mg/mL TCEP and 30 equivalents of PdCl_2_ were added and the pH was adjusted to 7. After stirring at 30 °C for 2 hours, a small amount of reaction solution was taken out and DTT was added with a final concentration of 500 mM. The progress of the Acm removal was meticulously monitored via analytic RP-HPLC (XB-C18 column, Welch).

A one-pot deprotection and ligation strategy was employed. Following the complete removal of the Acm protecting group, 4-mercaptophenyl acetic acid (MPAA, 300 eq. relative to the H2AK119C-CAET-SH peptide) was added, and the pH was adjusted to 5.0 to dissolve MPAA. A 1.5-fold molar excess of Ub G75-MesNa was added to the reaction mixture, and the pH was adjusted to 6.5. After reacting at 30 °C for 16 hours, 600 equivalent of TCEP was added to quench the reaction. The final product of H2AK119C-CAET-Ub was purified by semi-preparative RP-HPLC (XB-C18 column, Welch) and lyophilized.

Lyophilized H2AK119C-CAET Ub was dissolved in activating buffer (100 mM HEPES, 6 M Gn·HCl, pH 7.8) at a final concentration of 0.3 mM. Then 27 equivalents of 2,2’-Dithiodipyridine (AT_2_, CAS: 2127-03-9, 1 M stock in DMF) was diluted with buffer (100 mM HEPES, 6 M Gn HCl, pH 7.8). H2AK119C-CAET-Ub was added dropwisely to the AT_2_ solution in a 30 °C water bath for 15 minutes. H2AK119C-CAET Ub AT powder was obtained through semi preparative HPLC (XB-C18 column, Welch) purification and freeze-drying.

### Chemical synthesis of fluorescein-labeled H2A (86-129)-K119Ub peptide (FITC-H2A-Ub)

The component H2A (A86C-129)-K119 auxiliary groups (αG) was synthesized via solid-phase peptide synthesis (SPSS, Liberity Blue 2.0, CEM), and ligated to Ub (1-75) hydrazide as previously described^49^. The final products H2A (A86C-129)-K119Ub was purified by RP-HPLC (XB-C18 column, Welch) and lyophilized. The fluorescein was labeled on the peptide by incubating the H2A (A86C-129)-K119Ub peptide (1 mM) with 5 mM fluorescein-5-isothiocyanate (CAS number: 27072-45-3) for 30 min at 37℃. Products were purified by HPLC (XB-C18 column, Welch) and verified using MS. Lyophilized FITC-H2A-Ub was dissolved in unfolding buffer (6 M Gn·HCl, 50 mM HEPES pH 7.5). Refolding was initiated by gradual addition of refolding buffer (50 mM HEPES pH 7.5, 150 mM NaCl) to reduce Gn·HCl to a final concentration of 0.5 M.The refolded FITC-H2A-Ub was purified by size-exclusion chromatography (Superdex 75 10/300 GL, Cytiva). Purified FITC-H2A-Ub was collected and stored at –80°C.

### Disulfide crosslinking reactions of H2AK119Ub^AT^ Probes with DUB

H2AK119Ub^AT^ nucleosome was mixed with BAP1/ASXL1 or BAP1/ASXL1^K351Ub^ at a stoichiometric ratio of 1:2.2 in crosslinking buffer (25 mM HEPES pH 7.5, 50 mM NaCl, 2 mM MgCl_2_, 20 μM ZnCl_2_), with a 10 μL final volume. The final concentration of H2AK119Ub^AT^nucleosome and DUB (BAP1/ASXL1 or BAP1/ASXL1^K351Ub^) were 13.6 μM and 30 μM, respectively. The reaction system was placed at 30 ℃ for 2 h, and the reaction process was monitored by Coomassie-stained SDS-PAGE. For the crosslink of H2AK119Ub^AT^ nucleosomes and USP16, a 1:2.5 molar ratio was used.

### AlphaLISA

The AlphaLISA assay was performed in 384-well white OptiPlate (Revvity) with a final reaction volume of 40 μL. 10 μL of catalytically inactive (BAP1 C91A) BAP1/ASXL1 or BAP1/ASXL1^K351Ub^ with varied concentration (50, 25, 12.5, 6.25, 3.13, 1.56, 0.78, 0.39, 0.20, 0.10 μM) were incubated with anti-6×His-acceptor beads (Perkin Elmer, Cat: 2947839) for 2 h at room temperature in the binding buffer (50 mM HEPES pH 7.5, 150 mM NaCl, 0.1% v/v Tween-20, 0.1% v/v BSA, 1 mM dithiothreitol). 10 μL of 25 nM (final concentration) biotinylated H2AK119Ub nucleosomes were incubated with streptavidin-donor beads (Perkin Elmer, Lot: 676002S) for 2 h at room temperature. The enzyme and nucleosome carried the beads were mixed and incubated for 30 min. The AlphaLISA signal was quantified using an EnVision Plate Reader (Perkin Elmer). Data were analyzed with GraphPad Prism X software. All incubation was performed in dark.

### Electrophoretic mobility shift assay (EMSA)

Catalytically inactive BAP1 (C91A) complexes (BAP1/ASXL1 or BAP1/ASXL1^K351Ub^) were prepared in a 12-point 1:1 serial dilution series (12.8, 6.4, 3.2, 1.6, 0.8, 0.4, 0.2, 0.1, 0.05, 0.025, 0.0125, 0.00625 μM) using binding buffer (50 mM HEPES pH 7.5, 150 mM NaCl, 0.1% BSA, 1 mM DTT), followed by incubation with 10 nM (final concentration) H2AK119Ub nucleosomes for 15 min on ice in a final volume of 4 μL. 1 μL 5×native loading buffer (0.25% m/v bromophenol blue dissolved in 50% glycerol) was mixed with the reaction buffer gently. Native-PAGE analysis was used to separate DUB-nucleosome complex and nucleosome, and detected by SYBR-Gold staining.

### Molecular Dynamic simulation

The cryo-EM structures of BAP1/ASXL1^K351Ub^ and BAP1/ASXL1 complexes with the H2AK119Ub nucleosome substrate computationally removed were used as the initial models for simulations. The protein structure was preprocessed using the Protein Preparation Workflow (PPW) at pH 7.4^52^. This preparation included addition of hydrogen atoms, formation of disulfide bonds, completion of missing side chains, capping of termini, and removal of crystallographic water molecules. The structure underwent all-atom energy minimization using the OPLS4 force field with a maximum RMSD constraint of 0.3 Å^53^.

The prepared structure was solvated in a TIP3P water model within an orthorhombic box with a 10 Å buffer distance. Physiological conditions were simulated by adding 0.15 M NaCl, with additional Na+ counterions included to neutralize the system. All simulations were conducted using the OPLS4 force field implemented in the Desmond MD system within the Schrödinger Suite (Release 2024-2).

The constructed systems underwent equilibration prior to production simulations, eliminating structural strain through energy minimization, followed by short molecular dynamics simulations with default parameters. NPT ensemble class (pressure = 1 bar; temperature = 300 K) was used. The relaxed systems were subjected to three independent 500 ns MD simulation production runs, each initialized with different velocity seeds to ensure thorough sampling of the conformational space. Trajectory and energy data were collected at 100 ps intervals, yielding 5,000 frames per production run for subsequent analysis.

Trajectory analysis was performed to calculate RMSD and RMSF values with reference to the initial frame. Seven critical residues (BAP1: E7, Y33, F168, I226, N229; ASXL1: E311, H315) that form the substrate ubiquitin-binding pocket were identified for detailed characterization. Structural rigidity and integrity of the substrate ubiquitin-binding pocket were assessed using the trajectory_asl_monitor.py, which tracked distances between critical residue groups throughout the simulation with a 20 Å cutoff threshold. Representative conformations of the substrate ubiquitin-binding pocket were identified through clustering analysis using trj_cluster.py, employing the centroid hierarchical algorithm with a 1.5 Å cutoff threshold.

### Cryo-EM sample preparation

Prior to cryo-EM sample preparation, BAP1/ASXL1^K351Ub^ or nucleosomes were equilibrated in non-reducing buffer (20 mM HEPES, pH 7.5, 150 mM NaCl, 1 mM DTT, 5% glycerol). BAP1/ASXL1^K351Ub^ incubated with H2AK119Ub^AT^ nucleosomes at a 2.1:1 molar ratio for 170 minutes to facilitate disulfide crosslinking. The reaction was monitored 4–12% SDS-PAGE. Then equal volumes of crosslinking buffer (25 mM HEPES, 50 mM NaCl, 0.25% glutaraldehyde) added into the cross-linked complex for 10 minutes at room temperature, crosslinking was quenched by 50 mM Tris buffer (pH=8.0). The sample was concentrated to less then 100 μL using a centrifugal filter (Merck, 100 kDa molecular weight cutoff) and purified by size-exclusion chromatography (SEC) on a Superdex 6 Increase 5/150 GL column (Cytiva) pre-equilibrated with 25 mM HEPES, 50 mM NaCl. Peak fractions were analyzed by 4–12% SDS-PAGE (see **Extended Data Fig. 3a–b** for biochemical validation). The major peak eluting was pooled and concentrated for plunging.

For cryo-EM grid preparation, 3.5 μL of the sample was applied to pre-glow-discharged Quantifoil Au 300-mesh grids and incubated for 60 seconds. Grids were blotted for 3.0 seconds at 8°C and 100% humidity in a Thermo Scientific Vitrobot Mark IV, using blotting forces of −1, 0, or 1. Following blotting, grids were rapidly vitrified in liquid ethane and stored in liquid nitrogen.

### Cryo-EM data collection and image processing

The cryo-EM data processing workflow pipeline initiated with the collection of 7,994 micrographs, which were subjected to motion correction to mitigate beam-induced sample drift. From these, 10,914,932 raw particles were identified through initial particle picking and extracted at bin4 (4× downsampling) to optimize computational efficiency. A preliminary round of two-dimensional (2D) classification was employed to discard particles associated with evident contaminants. Subsequent *ab initio* three-dimensional (3D) classification (T=4), performed on a representative subset of particles, yielded initial density maps corresponding to two distinct structural classes:: 1 and 2:1.

For the 1:1 dataset (1,525,599 particles), a region-specific mask targeting the BAP1/ASXL1^K351Ub^ complex was applied during no-alignment 3D classification (T=4). Particles within classes exhibiting BAP1/ASXL1^K351Ub^ density were isolated and re-extracted at bin2. Iterative cycles of 3D auto-refinement and no-alignment 3D classification (T=28) improved particle homogeneity, refining the dataset to 578,842 particles processed at bin1. Global refinement and postprocessing (B-factor = −40) produced an initial resolution of 3.34 Å, which was elevated to 3.02 Å (FSC = 0.143) after CTF refinement and particle polishing. A subsequent no-alignment 3D classification step excluded low-resolution particles, yielding a final subset of 270,505 particles. These underwent 3D auto-refinement (3.51 Å) and postprocessing with a globular mask to achieve a 3.12 Å map. Focused refinement (B-factor = −100) further resolved the BAP1/ASXL1^K351Ub^ region, generating a 3.60 Å local map. A composite map was reconstructed by integrating the 3.12 Å globular map, 3.60 Å focused map, and 3.51 Å auto-refinement map.

The 2:1 dataset (822,144 particles), displaying BAP1/ASXL1K351Ub density on both nucleosome disc faces, underwent 3D auto-refinement followed by no-alignment 3D classification (T=26) using a mask targeting weaker BAP1/ASXL1K351Ub densities. From this, 166,213 particles with well-defined BAP1/ASXL1K351Ub density were re-extracted at bin1 and subjected to C2-symmetry-imposed 3D auto-refinement, yielding a 3.75 Å map. A final round of classification retained 131,081 particles, which were auto-refined (3.70 Å, C2 symmetry) and postprocessed (sharpened with a B-factor of −20) to generate a 3.27 Å reconstruction.

All resolutions were calculated using the FSC = 0.143 gold-standard criterion. Processing workflows were implemented in RELION-3.1^54^, with schematic summaries provided in Extended Data Fig. 3. Structural visualizations were rendered using ChimeraX (v1.6.1).

### Model building, refinement and validation

Atomic models were constructed based on cryo-EM density maps corresponding to a 1:1 stoichiometric complex. The following maps were utilized for iterative model building: (1) a 3.02 Å global map sharpened with a B-factor of −40, (2) a 3.12 Å global map sharpened at B-factor −60, and (3) a 3.51 Å locally refined map (3D auto-refinement) that provided enhanced density for the BAP1 α-helix (residues 56–60) interacting with the H2A-H2B acidic patch. Additionally, a 3.60 Å focused map of the BAP1/ASXL1^K351Ub^ subcomplex was used to resolve critical interfaces. Initial templates for the nucleosome, Ub^ASXL-K351^, and BAP1/ASXL1/Ub^H2AK1^^19^ complex were derived from PDB entries 7XD1, 1UBQ, and 8H1T, respectively. These partial models were rigid-body fitted into the density maps using ChimeraX v1.6.1 and merged into a unified atomic model using WinCoot v0.8.2^55^. Manual adjustments included mutating histone H3 residues C96 and C110 to serine and reverting BAP1 residue S91 (from PDB 8H1T) to cysteine to match biochemical data. Regions lacking interpretable density were truncated. The model underwent iterative real-space refinement in Phenix v1.19.2^56^ with secondary structure restraints applied to each subunit. Convergence was achieved through multiple rounds of refinement, including geometry regularization, clash minimization, and Ramachandran outlier correction. The final model was validated using the Phenix comprehensive validation (cryo-EM) module, which assessed global map-model correlations, local residue fit, and stereochemical accuracy. Key validation metrics, including Fourier shell correlation (FSC) curves, model-to-map cross-correlation values, and clash scores, etc. are summarized in **Extended Data Table 1**.

### Hydrogen-Deuterium Exchange Mass Spectrometry (HDX-MS)

Hydrogen-deuterium exchange was initiated by adding 45 μL deuterium buffer (50 mM HEPES pH 7.1, 150 mM NaCl, 1 mM DTT, 99% D2O) to 5 μL enzyme samples (BAP1/ASXL1 or BAP1/ASXL1K351Ub, 50 μM) at 25 ℃. Reactions were quenches at 30, 90, 300, or 1800 s by adding 50 μL ice-cold quenching buffer (4 M guanidine HCl, 200 mM citric acid, and 500 mM TCP, 100% H2O, pH 1.8). The samples were consequently placed on ice and 5 μL pepsin solution (1 μM) was added. After 3 min digestion, samples were placed in the Thermo-Dionex Ultimate 3000 HPLC system autosampler for data analysis. Deuterium uptake was quantified by centroid mass changes characterized using HDExaminer (version PD 1.4, Thermo Scientific).

## Data availability

The cryo-EM maps and atomic model of BAP1/ASXL1^K351Ub^ in complex with H2AK119Ub nucleosomes have been deposited in the Electron Microscopy Data Bank (EMDB) under accession codes: EMD-63884 (BAP1/ASXL1^K351Ub^/H2AK119Ub NCP complex in a 1:1 stoichiometry), EMD-63885 (BAP1/ASXL1^K351Ub^/H2AK119Ub NCP complex in a 2:1 stoichiometry), respectively. The atomic model of BAP1/ASXL1^K351Ub^ in complex with H2AK119Ub nucleosomes with a 1:1 stoichiometry has been deposited in the Protein Data Bank (PDB) under accession codes 9U5U. The ASXL1 K351 ubiquitination information was found in PhosphoSitePlus database (https://www.phosphosite.org/). Instrument schematic in **Extended Data Fig. 8a** was created with BioArt resources (BioArt: NIAID Visual & Medical Arts; 2025/2/11; bioart.niaid.nih.gov/bioart/579).

